# ExploreASL: an image processing pipeline for multi-center ASL perfusion MRI studies

**DOI:** 10.1101/845842

**Authors:** Henri Mutsaerts, Jan Petr, Paul Groot, Pieter Vandemaele, Silvia Ingala, Andrew D Robertson, Lena Václavů, Inge Groote, Hugo Kuijf, Fernando Zelaya, Owen O’Daly, Saima Hilal, Alle Meije Wink, Ilse Kant, Matthan W.A. Caan, Catherine Morgan, Jeroen de Bresser, Elisabeth Lysvik, Anouk Schrantee, Astrid Bjørnebekk, Patricia Clement, Zahra Shirzadi, Joost P.A. Kuijer, Udunna C. Anazodo, Dasja Pajkrt, Edo Richard, Reinoud P.H. Bokkers, Liesbeth Reneman, Mario Masellis, Matthias Günther, Bradley J. MacIntosh, Eric Achten, Michael A. Chappell, Matthias J.P. van Osch, Xavier Golay, David L. Thomas, Enrico de Vita, Atle Bjørnerud, Aart Nederveen, Jeroen Hendrikse, Iris Asllani, Frederik Barkhof

**Author notes:** authors contributed equally to this work (Henri Mutsaerts, Jan Petr). Corresponding author: Henri JMM Mutsaerts, MD PhD, @, Dep. of Radiology and Nuclear Medicine, PK -1, De Boelelaan 1117, 1081 HV Amsterdam, The Netherlands.

## Abstract

Arterial spin labeling (ASL) has undergone significant development since its inception, with a focus on improving standardization and reproducibility of its acquisition and quantification. In a community-wide effort towards robust and reproducible clinical ASL image processing, we developed the software package ExploreASL, allowing standardized analyses across centers and scanners.

The procedures used in ExploreASL capitalize on published image processing advancements and address the challenges of multi-center datasets with scanner-specific processing and artifact reduction to limit patient exclusion. ExploreASL is self-contained, written in MATLAB and based on Statistical Parameter Mapping (SPM) and runs on multiple operating systems. The toolbox adheres to previously defined international standards for data structure, provenance, and best analysis practice.

ExploreASL was iteratively refined and tested in the analysis of >10,000 ASL scans using different pulse-sequences in a variety of clinical populations, resulting in four processing modules: Import, Structural, ASL, and Population that perform tasks, respectively, for data curation, structural and ASL image processing and quality control, and finally preparing the results for statistical analyses on both single-subject and group level. We illustrate ExploreASL processing results from three cohorts: perinatally HIV-infected children, healthy adults, and elderly at risk for neurodegenerative disease. We show the reproducibility for each cohort when processed at different centers with different operating systems and MATLAB versions, and its effects on the quantification of gray matter cerebral blood flow.

ExploreASL facilitates the standardization of image processing and quality control, allowing the pooling of cohorts to increase statistical power and discover between-group perfusion differences. Ultimately, this workflow may advance ASL for wider adoption in clinical studies, trials, and practice.

## Introduction

Arterial spin labeling (ASL) is a non-invasive magnetic resonance imaging (MRI) technique with the potential of providing absolute quantification of cerebral perfusion *in vivo*. Since its inception almost three decades ago, ASL-based perfusion imaging has been increasingly used in basic neuroscience and clinical studies. The first decade of research following the invention of ASL in 1990 (Detre et al. 1992) consisted mainly of technical developments, such as the prolongation of the post-labeling delay in 1996 (Alsop and Detre 1996), background suppression in 1999 (Alsop and Detre 1999; Ye et al. 2000), and pseudo-continuous labeling in 2005 (Dai et al. 2008) – all features geared toward improving the signal-to-noise (SNR) ratio of ASL images. These advances enabled proof-of-principle studies using small clinical datasets, such as patients with cerebrovascular and neurodegenerative diseases (Detre et al. 1998; Alsop et al. 2000), epilepsy (Liu et al. 2001), brain tumors (Warmuth et al. 2003), as well as pharmacological applications (Wang et al. 2011; MacIntosh et al. 2008).

The second decade was primarily focused on the validation of the technique by way of clinical implementation (Deibler et al. 2008), evaluation of multi-center reproducibility (Petersen et al. 2010; Mutsaerts et al. 2015), and comparison with [^15^O]-H_2_O positron emission tomography (PET) (Heijtel et al. 2014). Several reproducibility studies showed that conventional ASL techniques had developed to the point where the intrinsic variance of the acquisition itself (Chen et al. 2011; Gevers et al. 2011; Heijtel et al. 2014; Mutsaerts et al. 2014) was close to or below physiological variance of perfusion (Joris et al. 2018; Clement et al. 2017). Another pivotal aspect for the use of ASL were pharmaceutical studies (Handley et al. 2013; MacIntosh et al. 2008).

Currently in its third decade, following the consensus recommendations for the acquisition and quantification of ASL images (Alsop et al. 2015), ASL is ready for large multi-center observational studies and clinical trials (Jack et al. 2010; Almeida et al. 2018; Blokhuis et al. 2017). However, despite the consensus in clinical implementation and image acquisition (Alsop et al. 2015), ASL image processing (Wang et al. 2008; Shin et al. 2016; Melbourne et al. 2016; Chappell et al. 2010; Mato Abad et al. 2016; Li et al. 2019; Bron et al. 2014) remains disparate among research laboratories. In literature, detailed description of all processing steps is often lacking. Clinical studies are often performed without proper quality control (QC) or with arbitrary QC metrics. This hampers both the interpretation and reproducibility of individual studies as well as meta-analyses of multiple studies. A consensus on the best practices to robustly process ASL data would facilitate comparison of results across centers and studies, avoid duplicate development, and speed up the translation into clinical practice, as is advocated by the Open Source Initiative for Perfusion Imaging (OSIPI) (www.osipi.org).

For these reasons, the software package ExploreASL was initiated through the EU-funded ASL workgroup COST-action BM1103 “ASL In Dementia” (www.aslindementia.org) with the aim of developing a comprehensive pipeline for reproducible multi-center ASL image processing. To date, ExploreASL has been used in more than 30 studies consisting of more than 10,000 ASL scans from three MRI vendors - GE, Philips, Siemens, with pulsed ASL and pseudo-continuous ASL (PCASL) sequences (Mutsaerts et al. 2019). The primary aims of ExploreASL are to increase the comparability and enable pooling of multi-center ASL datasets, as well as to encourage and facilitate cross-pollination between clinical investigators and image processing method developers.

## Theory: Software Overview

ExploreASL is developed in MATLAB (MathWorks, MA, USA, tested with versions 2011-2019) and uses Statistical Parametric Mapping 12 routines (SPM12, version 7219) (Ashburner et al. 2012; Flandin et al. 2008). Here, we describe the implementation of ExploreASL version 1.0.0, which is available as a compiled version with a manual on www.ExploreASL.org. ExploreASL provides a fully automated pipeline that comprises all the necessary steps from data import and structural image processing to cerebral blood flow (CBF) quantification and statistical analyses. Unique features of ExploreASL include:

- self-contained software suite: all third-party toolboxes are included in the installation, compatible with Linux, macOS, and Windows and supporting multi-threading;
- flexible data import from different formats including (enhanced) DICOM, Siemens’ MOSAIC variant, Philips PAR/REC, NIfTI and Brain Imaging Data Structure (BIDS) (Gorgolewski et al. 2016), with automatic detection of control-label or label-control order;
- data management: anonymization, compression of image files;
- modular design: automatically iterates over all available subjects and scans, allows to suspend and resume processing at any point, allows investigators to change/replace each sub-module;
- image processing optimized for: multiple centers, different ASL implementations from GE/Philips/Siemens (Mutsaerts et al. 2018), both native/standard-space analysis, advanced ASL markers ― e.g. spatial coefficient-of-variation (CoV) (Mutsaerts et al. 2017), asymmetry index (Kurth et al. 2015), and partial volume correction (PVC) (Asllani et al. 2008);
- extensive QC and data provenance: visual QC for all intermediate and final images, comparison with perfusion templates from different ASL implementations, progress report with processing history (provenance).

In the following sections, we review each processing step of the four ExploreASL modules as outlined in Figure 1 and summarized in Table 1. Each section starts with a brief methodological review including the rationale within the context of ASL processing, followed by a detailed description of the ExploreASL implementation, and ending with a discussion of emerging developments and potential future improvements.

**Figure 1.**
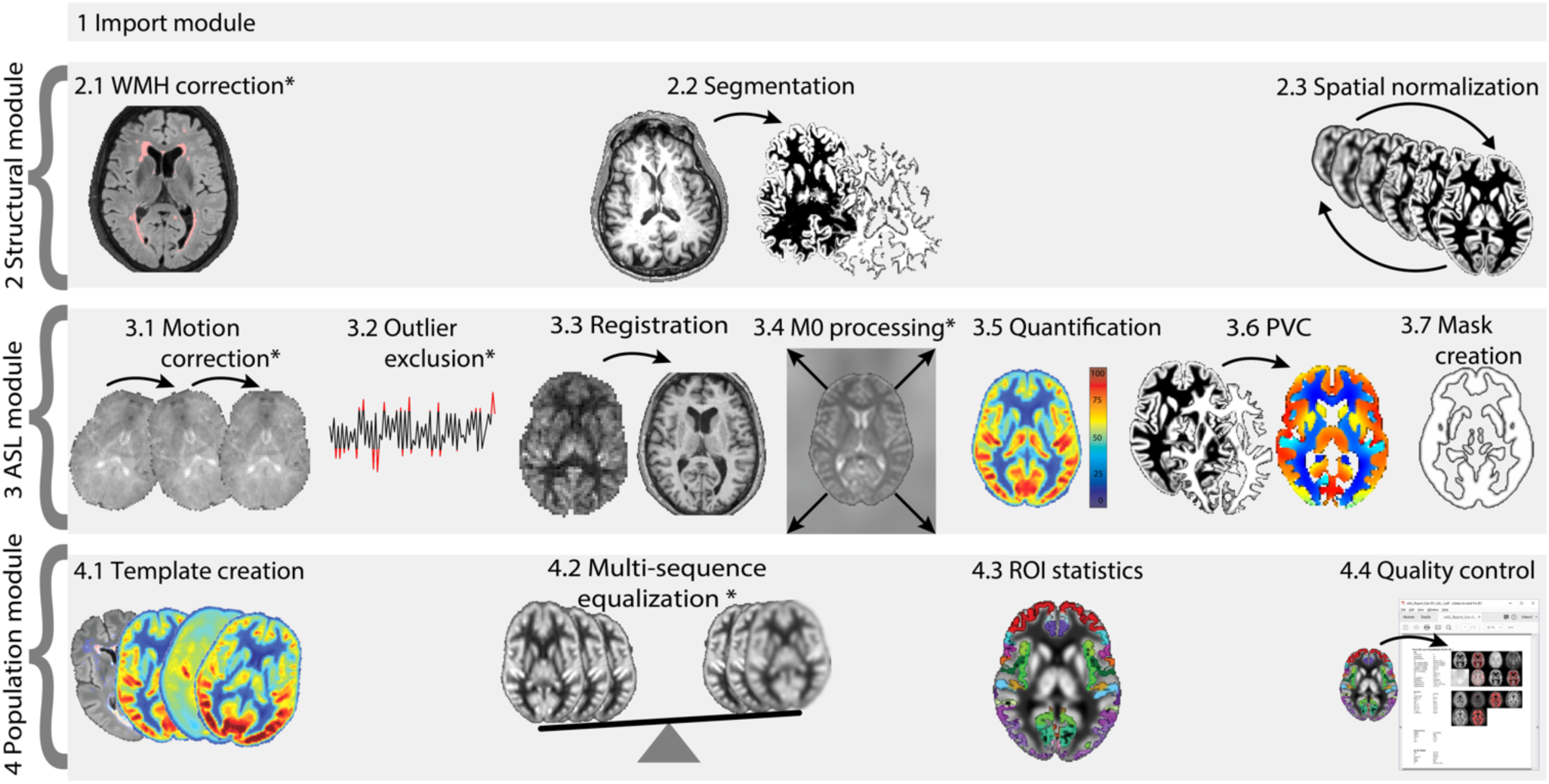
Schematic diagram of ExploreASL processing steps. Steps marked with a * are optional, e.g. when FLAIR, ASL time-series, or M0 scans are available. PVC = partial volume correction, ROI = regions of interest, WMH = white matter hyperintensity. The population module can be run on a single subject level, as well as on one or multiple populations/centers/cohorts or other groups.

**Table 1.** Overview of image processing steps and implementation in ExploreASL. ASL = arterial spin labelling, BIDS = Brain Imaging Data Structure, CAT = Computational Anatomic Toolbox, CBF = cerebral blood flow, CoV = coefficient of variation, ΔM = perfusion-weighted difference image, dcm2niiX (Li et al. 2016), DICOM = Digital Imaging and COmmunications in Medicine, ENABLE = ENhancement of Automated BLood flow Estimates, FoV = field-of-view, LGA = Lesion Growth Algorithm, LPA = Lesion Prediction Algorithm, LST = Lesion Segmentation Toolbox, MNI = Montreal Neurological Institute, NIfTI = NeuroImaging Informatics Technology Initiative, QC = quality control, pGM = gray matter partial volume, PLD = post-labeling delay, PSF = point spread function, PVC = partial volume correction, r = release, ROI = region of interest, SD = standard deviation, SNR = signal-to-noise ratio, SPM = Statistical Parametric Mapping, tSNR = temporal SNR, tsv = tab-separated value, WMH = white matter hyperintensity, Zig-zag = “zig-zag” regressor.

| Processing step | ExploreASL implementation | Specifics, optional features |
| --- | --- | --- |
| **1. Import module** |  |  |
| 1.1 Data import | dcm2niiX | Converts DICOM to NIfTI, supports MOSAIC, PAR/REC, BIDS |
| **2. Structural module** |  |  |
| 2.1 WMH correction | LST2 (r2.0.15) - LPA (LGA optional) | Segments WMH and fills lesions on T1w; improves T1w segmentation |
| 2.2 Segmentation | CAT12 (r1363) | Outputs partial volume maps; supports lesion cost function masking |
| 2.3 Spatial normalization | Geodesic Shooting | Uses CAT12 template, supports creation of study-specific templates |
| **3. ASL module** |  |  |
| 3.1 Motion correction | SPM12 realign | Realigns ASL control/label images to mean position, uses a zig-zag control-label regressor |
| 3.2 Outlier exclusion | ENABLE | Removes motion peaks, uses tSNR optimization |
| 3.3 Registration | SPM12 rigid-body | Registers $\Delta M$-pGM or M0-T1w |
| 3.4 M0 processing | M0 image | Masks, smooths, extrapolates M0 to avoid division artifacts |
| 3.5 CBF quantification | Consensus paper model | Computes CBF based on the single compartment model, single PLD; supports dual compartment |
| 3.6 PVC | Linear regression | Performs PVC on kernel or ROI basis, optionally estimates the effective spatial resolution/PSF of ASL |
| 3.7 Analysis mask creation | Combine individual masks | Combines FoV, susceptibility artifacts and vascular artifacts, use $p > 0.95$ of population masks |
| **4. Population module** |  |  |
| 4.1 Template creation | Final images | Calculates population mean, SD, CoV, SNR, also for intermediate images |
| 4.2 Multi-sequence equalization | Remove residual sequence-specific effects | Equalizes bias fields, spatial CoV, and smoothness, uses sequence-specific templates |
| 4.3 ROI statistics | CBF and spatial CoV, with or without PVC | Uses MNI structural, Harvard-Oxford, Hammers, and custom atlases |
| 4.4 Quality control | Single-subject PDF report | Performs QC of images, DICOM values, volumetrics, motion etc. Outputs population report in TSV files |

## Theory: Implementation

### 1. Import module

To avoid manual restructuring of arbitrary data structures from the scanner or other sources (Fallis et al. 2013), ExploreASL uses a flexible input data/directory description scheme based on regular expressions and converts the data to a BIDS-compatible data structure (Gorgolewski et al. 2016); the full BIDS ASL extension is currently in development (bids.neuroimaging.io). The input images can be NIfTI format, conventional or enhanced DICOM, Philips PAR/REC, or Siemens mosaic format, which are then converted to NIfTI using dcm2niiX (Li et al. 2016). ASL images can be provided as control-label time-series, a single perfusion-weighted image, or an already quantified CBF image, from any 2D or 3D readout schemes, and from any MRI vendor. Before an image is processed, ExploreASL first computes and aligns the center-of-mass of each image to the origin of the world coordinates to deal with potentially incorrectly stored orientations. Additionally, ExploreASL provides an overview of missing and unprocessed files, automatically detects the order of control and label images from the image intensities, and checks the DICOM tags of repetition and echo time and scale factors/slopes across individuals.

### 2. Structural module

This module process the structural images by the following steps: 2.1) segments the white matter (WM) hyperintensities (WMH) on fluid-attenuated inversion recovery (FLAIR) images and uses them to fill the corresponding WM hypointensities on the T1-weighted (T1w) images., 2.2) the structural images are subsequently segmented into gray matter (GM), white matter, and cerebrospinal fluid (CSF) maps, and 2.3) normalized to the MNI standard space (Evans et al. 2012). The segmentations are used to obtain tissue partial volume (PV) fractions for computation of CBF (Asllani et al. 2008). The registration transformations are used to bring ASL images acquired from different sessions and/or different subjects in the same space and thus facilitate visual comparison in the same space, automatic QC, as well as group analysis.

#### 2.1. WMH correction

The presence of WMH can affect the GM/WM classification of T1w images in two ways: 1) WMH themselves can be incorrectly segmented as GM; 2) image intensities of WMH bias global modeling of GM and WM intensity distributions (Pareto et al. 2016; Battaglini et al. 2012). ExploreASL alleviates these complications by lesion-filling the T1w image before initiating the segmentation (Battaglini et al. 2012): voxel intensities in the hypointense WMH regions on the T1w images are replaced by bias field-corrected values from the surrounding, normal-appearing WM (Chard et al. 2010) (Figure 1). The Lesion Segmentation Toolbox (LST, version 2.0.15) is used because of its empirically proven robustness, scanner independence, and non-reliance on the requirement of a training set (de Sitter et al. 2017). LST detects outliers in the FLAIR WM intensity distribution and assesses their likelihood of being WMH (Schmidt et al. 2012). While ExploreASL offers the option of both LST lesion growing and lesion prediction algorithms, the default is set to the latter, which has been shown to be more robust (de Sitter et al. 2017). This WHM correction described here is only performed when FLAIR images are available. Optionally, the WMH segmentation can be skipped by providing an external WMH segmentation; then only the LST clean-up procedure is performed to remove classification errors from this external input (Schmidt et al. 2012).

#### 2.2. Segmentation

To segment the 3 main tissue classes GM, WM, and CSF, ExploreASL uses the Computational Anatomy Toolbox 12, release 1363 (CAT12, the successor of VBM8) (Gaser et al. 2009) for SPM12. CAT12 allows local variations in the tissue intensity distributions, making it more robust to the presence of pathology such as tumors, edema, and WM lesions (Battaglini et al. 2012; Petr et al. 2018) (Supplementary Figure 1). CAT12 works on all operating systems (OS), and has been shown to outperform other available methods such as FreeSurfer v5.3.0, FSL v5.0, and SPM12 (Mendrik et al. 2015). The CAT12 segmentation algorithm is based on improvements of Unified Segmentation (Ashburner et al. 2005), two essential improvements being that it allows spatially varying GM-WM intensity distributions, and provides PV maps rather than posterior probability maps (Tohka et al. 2004).

#### 2.3. Spatial normalization

For non-linear registration to MNI space ExploreASL uses Geodesic Shooting (Ashburner et al., 2011) - the successor of Diffeomorphic Anatomical RegisTration using Exponentiated Lie algebra (DARTEL) (Ashburner et al. 2007) - within the CAT12 toolbox (Gaser et al. 2009). The reason for this choice is that CAT12 has a single subject implementation using the IXI adult template, brain-development.org/ixi-dataset. Optionally, new templates can be created by these SPM toolboxes on a population level, e.g. for populations where an adult template is not sufficient. Although alternative methods (Klein et al. 2010) may outperform DARTEL/GS in specific populations, the default settings of DARTEL and Geodesic Shooting are sufficiently tested in clinical studies to provide adequate performance across different populations and scanners (Ripollés et al. 2012). We adapted the CAT12 segmentation algorithm to offer the possibility to input customized segmentations of structural lesions such as space-occupying lesions or cerebral infarcts such that the lesion region is ignored by the non-linear registration (Crinion et al. 2007) (Supplementary Figure 2). To avoid unrealistic deformations within the lesion, the non-linear transformation is only used outside the lesion; whereas the initial SPM transformation (low degree-of-freedom non-linear) is used within the lesion. These two transformation fields are combined with a gradual transition margin around the lesion amounting to 10% of the lesion volume.

ExploreASL offers the option to register longitudinal ASL studies with the SPM12 module for longitudinal registration (Ashburner and Ridgway 2012), which takes the similarity between structural images from the same subjects into account. The first time point is used as a reference for both within- and between-subject registration. However, this requires further validation in the presence of large brain deformations between sessions, such as tumors, resections, or infarcts (Petr et al. 2018).

### 3. ASL module

This module processes the ASL images by 3.1) correcting for motion, 3.2) removing outliers, 3.3) registering with the structural data, and by 3.4) processing the M0 images. Then, 3.5) the CBF is quantified with correction for hematocrit and vascular artifacts, after which 3.6) the PV effects are corrected for. All image processing described below is performed in native space, unless stated otherwise. All intermediate and final images are also transformed into standard space for QC and group analyses.

#### 3.1. Motion correction

The adverse effects of head motion can be partly alleviated by correcting for motion using image processing (Alsop et al. 2015). Traditionally, head motion is estimated assuming a 3D rigid-body transformation with a sum-of-squares cost function (Wang et al. 2008; Mato Abad et al. 2016). However, because the average control-label intensity difference can be partly interpreted by the algorithm as motion, some investigators perform motion estimation separately for the control and labeled images (Wang et al. 2008). Instead, in ExploreASL, an adaptation of the SPM12 motion correction is used, which minimizes apparent motion attributable to the control-label intensity difference from the estimated motion parameters using a “zig-zag” regressor (Wang et al. 2012) (Supplementary Figure 3).

#### 3.2. Outlier exclusion

Despite motion correction, large motion spikes can still have a significant negative effect on the ASL image quality, especially when they occur between control and label images (Wang et al. 2008). In fMRI literature, peak motion relative to mean individual motion is often excluded based on a set threshold, e.g. RMS of 0.5 of the voxel size (Power et al. 2012). ExploreASL uses a threshold-free method named ENhancement of Automated BLood flow Estimates (ENABLE) (Shirzadi et al. 2015), which sorts control-label pairs by motion and cumulatively averages them until the addition of further pairs significantly decreases the temporal voxel-wise signal stability (Supplementary Figure 4). The ExploreASL implementation of ENABLE employs the median GM voxel-wise temporal SNR (tSNR) as the criterion for signal stability (Shirzadi et al. 2018), regularized by an empirically-defined minimum tSNR improvement of 5%. ENABLE can also remove non-motion-related outliers, since other acquisition artifacts can be picked up by the motion estimation algorithm (Supplementary Figure 4).

In addition to motion, ASL images can contain a variety of acquisition and physiological artifacts including fat-shift artifacts, RF instability, gradient amplifier failure, labeling instability, and blood pulsatility artifacts. Several correction algorithms using ASL time series have been proposed which typically exclude voxels, slices or volumes as outliers, based on temporal and/or spatial signal distribution of the individual pairwise subtracted images (Bibic et al. 2010; Spann et al. 2017; Maumet et al. 2014; Tan et al. 2009; Dolui et al. 2017). Their applicability needs validation, however, as they base their correction criteria on the same parameter that is investigated (i.e. CBF), and/or they do not account for GM-WM perfusion differences. ExploreASL currently relies on the fact that ENABLE (Shirzadi et al. 2018) also partly removes outliers, as it operates relatively independent of (patho-)physiological changes of the signal intensity in the pairwise subtracted images (Robertson et al. 2017; Li et al. 2018).

#### 3.3. Registration

Accurate registration between the ASL and structural space is a critical step as registration errors are propagated to subsequent stages and analyses of CBF data. Specifically, the relatively large CBF differences between GM, WM, and CSF, mean that small misalignments can have a large impact on the accuracy of tissue-specific CBF quantification (Mutsaerts et al. 2018).

The image registration steps implemented in ExploreASL are based on a previous study in which the performance of several registration options were compared (Mutsaerts et al. 2018). Briefly, the registration of ΔM to gray matter partial volume (pGM) outperformed the registration of M0 to T1w, except for cases where the ΔM contrast was dissimilar to the pGM contrast (e.g. vascular artifacts, labeling artifacts, perfusion pathology). Rigid-body transformation proved to be a robust default choice (Mutsaerts et al. 2018), especially in the presence of pathology (Wang et al. 2008; Macintosh et al. 2010). Therefore, ExploreASL performs a ΔM-pGM rigid-body registration by default. The M0-T1w is used instead when macrovascular signal predominates tissue signal (spatial CoV above 0.6) (Mutsaerts et al. 2018). Note that such images are typically excluded from CBF statistics and only included when analyzing vascular parameters, such as the spatial CoV. For future validation, ExploreASL offers the option to register to the atlas of spatially normalized mean M0 images, CBF images, or CBF images with a high number of vascular artifacts created for different vendors and ASL sequences from previously processed large ASL datasets.

The rigid-body transformation does not account for the geometric distortion typical for 2D echo-planar imaging (EPI) or 3D GRadient And Spin Echo (3D GRASE) ASL images (Gai et al. 2017). Such deformations can be partially corrected with B0 field maps or M0 images with reversed phase-encoding direction (Madai et al. 2016) - which is implemented as option in ExploreASL by calling FSL TopUp (Andersson et al. 2003). Affine and uniform non-linear transformations, such as FNIRT or SPM’s ‘unified segmentation’ (Klein et al. 2009) can outperform the rigid-body transformation in the ΔM-pGM registration (Petr et al. 2018), although this remains to be validated in the presence of pathology.

#### 3.4. M0 processing

The measured control-label magnetization difference is proportional to the equilibrium magnetization (M0) of blood. Ideally, blood M0 would be measured in voxels containing only arterial blood, but that is not usually possible due to the relatively low spatial resolution of ASL images. Instead, M0 is calculated from either the brain tissue or CSF signal intensity (Çavuşoğlu et al. 2009). The use of the tissue-based M0 is recommended (Alsop et al. 2015) because of its ability to account for acquisition-specific effects such as variations in receive coil inhomogeneity or T2(*) weighting. For these reasons, ExploreASL by default processes an M0 image, and optionally supports the use of a single CSF M0 value (Çavuşoğlu et al. 2009; Pinto et al. 2019).

Currently, no consensus exists on whether the M0 should be quantified separately for GM and WM tissue types, especially for longer repetition time with a distinct GM-WM contrast. The M0 quantification can potentially be improved by using tissue specific quantification parameters - such as blood-brain partition coefficients λ and tissue relaxation times (Çavuşoğlu et al. 2009), and/or partial volume correction (Ahlgren et al. 2018). However, this can induce quantification errors in cases of suboptimal ASL-M0 registration.

ExploreASL aims to deliver consistent M0 quantification for multi-center populations with M0-scans acquired at different repetition time and different effective resolutions. ExploreASL smooths the M0 image with a large kernel (Beaumont 2015) after it has been masked for WM (Supplementary Figures 5-6) and rescaled to the mean GM M0 to account for B1 differences between GM and WM. This approach reduces the M0 image into a smooth bias field with the same smoothness/effective resolution for all ASL sequences and participants, and optimal SNR, while still canceling out acquisition-specific B1-field related intensity inhomogeneity. This makes the M0 image more robust and less sensitive to misalignment, and thus more consistent between ASL sequences (Mutsaerts et al. 2018) and individuals (Deibler et al. 2008). ExploreASL has the option to additionally mask the M0 bias-field for lesions that affect the M0 - e.g. brain tumors - and interpolate the M0 signal from the relatively unaffected brain regions (Croal et al. 2019).

#### 3.5. CBF quantification

An in-depth overview of ASL CBF quantification has been provided previously (Alsop et al. 2015; Chappell et al. 2018). ExploreASL uses the previously recommended single compartment quantification approach for clinical studies (Alsop et al. 2015), with options to use the dual compartment model, and/or to provide the hematocrit or blood T1 values. The previously recommended single compartment model assumes that the label decays with arterial blood T1 only (Alsop et al. 2015). Although a two-compartment model can provide CBF values that are in closer agreement with [^15^O]-H_2_O PET (Heijtel et al. 2014), this is often not feasible when blood T1, tissue T1, and micro-vascular arterial transit time are unknown, or would result in a constant scaling factor when assuming literature values. For these reasons, ExploreASL uses the single compartment model by default, and offers the two-compartment model as an optional feature.

The ASL label relaxes with the T1 of blood, a parameter that depends on hematocrit (Hales et al. 2016). Not taking hematocrit or blood T1 into account can lead up to 10-20% CBF overestimation for hematocrit as low as 17% (Vaclavu et al. 2016). Accounting for hematocrit is particularly relevant for between-group or longitudinal hematocrit changes e.g. due to treatment, which can be expected in certain populations or diseases (De Vis et al. 2014). ExploreASL allows to adjust for individual arterial blood T1 by either providing its value directly (Li et al. 2017) or by providing the hematocrit value and computing the blood T1 (Hales et al. 2016). As hematocrit and blood T1 measurements can be noisy - especially when obtained at different laboratories - a pragmatic approach is to apply the average blood T1 correction on a population rather than on an individual level (Elvsåshagen et al. 2018). Additionally, hematocrit and blood T1 can be modeled based on age and sex (Hales et al. 2014), but this requires validation. Note that after correcting the above-mentioned methodological effect, hematocrit might be still associated with CBF physiologically: hematocrit decreases or increases leading to compensatory hyper- or hypoperfusion.

#### 3.6. Partial volume correction

Since the spatial resolution of ASL is relatively low, a typical ASL voxel contains a mixture of GM, WM, and CSF signal, which is referred to as the partial volume effects. As the GM-WM CBF ratio is reported to lie between 2 and 7 (Asllani et al. 2008; Pohmann 2010; Zhang et al. 2014; Law et al. 2000), the tissue partial volume in each voxel has a large influence on the ASL measurement (Supplementary Figure 7). For these reasons, PVC (Asllani et al. 2009) is essential in studies that aim to differentiate structural changes (e.g. atrophy) from perfusion changes (e.g. related to neurovascular coupling) (Steketee et al. 2016). Several PVC algorithms have been proposed (Chappell et al. 2010; Zhao et al. 2017; Asllani et al. 2008; Liang et al. 2013), which assume locally homogeneous GM and WM CBF. Instead in some studies, GM volume is used as a covariate in the statistical analysis (Chen et al. 2011). Note that while PVC, in theory, corrects only for the PV effects and takes into account the intra- and inter-subject variability of the GM-WM CBF ratio, GM covariation can additionally affect the estimated physiological correlation between GM CBF and GM volume (Petr et al. 2018).

ExploreASL employs two versions of PVC, both based upon the most frequently used PVC, i.e. linear regression (Asllani et al. 2008): 1) a 3D Gaussian instead of a 2D flat kernel (default, referred to as “voxel-wise”) (Oliver 2015), or 2) computing PV-corrected CBF within each anatomical or functional region of interest (ROI) separately instead of using a kernel. Whereas the voxel-wise option allows further voxel-based analysis (VBA), the ROI-based PVC is in theory beneficial for a ROI-based analysis as effectively the kernel-size is selected based upon the anatomical ROI, which should be less sensitive to local segmentation errors. Moreover, it avoids cross-talk between ROIs. It still needs to be investigated how to define regions of optimal shape with respect to PVC performance, which depends on the spatial uniformity and SNR of the GM and WM CBF, and partial volume distributions within the ROI. To evaluate the effects of PVC, ExploreASL exports CBF maps and ROI values both with and without PVC.

For proper PVC or ROI definition, the true acquisition resolution - which often differs from the reconstructed voxel size - needs to be taken into account (Petr et al. 2018). This is especially important for 3D readouts, where the through-plane PSF can be up to 1.9 times the nominal voxel-size (Vidorreta et al. 2013; Vidorreta et al. 2014). Effects such as motion (Petr et al. 2016) and scanner reconstruction filters can contribute to further widening of the PSF of the final image. ExploreASL by default uses previously estimated true acquisition resolutions (Vidorreta et al. 2013; Vidorreta et al. 2014; Petr et al. 2018) and can optionally perform a data-driven spatial resolution estimation (Petr et al. 2018) that is generalizable to all ASL acquisitions. Contrary to alternative PSF estimations based on temporal noise autocorrelation (Cox 2012) or simulations of the acquisition PSF (Vidorreta et al. 2013; Vidorreta et al. 2014), this method does not require time series and inherently accounts for other sources of blurring (e.g. smoothing by motion and/or image processing) and is applicable without having detailed information about the sequence parameters needed to calculate the resolution from the k-space trajectory. However, this method requires further validation, especially in the presence of ASL image artifacts.

Lastly, the GM/WM maps obtained from the high-resolution structural images need to be downsampled to the ASL resolution before they are used for PVC or for ROI delineation in native space. A trivial interpolation to lower resolution may introduce aliasing, which can be addressed by applying a Gaussian filter - or a convolution with the PSF, if the PSF is known - prior to downsampling (Cardoso et al. 2015). It is important to note that the ASL image often has an anisotropic resolution and may be acquired at a different orientation compared to the structural image. To correct for this effect, ExploreASL pre-smooths the structural images with a Gaussian kernel of which the covariance matrix takes the orientation and PSF differences between the ASL and structural images into account (Cardoso et al. 2015).

#### 3.7. Analysis mask creation

For the statistics performed in section 4.3 - as well as for any voxel-based group statistics - an analysis mask aims to exclude voxels outside the brain or voxels with artifactual signal (e.g. macro-vascular, signal dropout) and restrict the analysis to regions with sufficient SNR and/or statistical power. This also avoids over-penalizing statistical power by family-wise error corrections. The susceptibility and field-of-view (FoV) masks are combined in section 4.3 into a group mask. The vascular masks are applied subject-wise to reflect the individual differences in vascular anatomy.

First, regions outside of the ASL FoV are identified, as whole brain coverage is not always achieved (Supplementary Figure 8). Second, a mask is created to remove voxels with intravascular signal. While intra-vascular signal - resulting from an incomplete tissue arrival of labeled spins - can be clinically useful (Mutsaerts et al. 2017; Mulhollan et al. 2018), it biases regional CBF estimates. The relatively large local temporal variability of such vascular artifacts can be detected in time series, in multi-post-labeling delay (PLD) acquisitions (Chappell et al. 2010) or by an independent component analysis (ICA) (Hao et al. 2018). ExploreASL uses a vascular artifact detection approach that is suitable for both single and multi-PLD ASL images. It identifies clusters of negative apparent CBF (Maumet et al. 2012) and voxels with extreme positive apparent CBF (Supplementary Figure 9). One potential caveat of masking out vascular voxels is the violation of the stationarity criterion of parametric voxel-wise statistics. While excluding voxels with high signal can violate the stationarity criterion of the ASL signal, there is currently no validated method that would be able to reliable estimate the perfusion and vascular signal contribution in such voxels from single-PLD data.

Second, regions with susceptibility signal-dropout artifacts are removed. Regions frequently having low SNR for 2D EPI and 3D GRASE ASL include the orbitofrontal cortex near the nasal sinus and the inferior-medial temporal gyrus near the mastoid air cavities. There are several methods that we decided not to include in ExploreASL: thresholding the M0 or the mean control image to identify signal dropout (Wang 2014) or masking them with FSL BET (Smith 2002), as this may fail with background suppression, blurred 3D acquisitions, poor ASL-M0 registration, or a strong bias field (Mutsaerts et al. 2018). If multiple individual unsubtracted control-label images are available, a mask could be created based on the tSNR of the control or label images. However, time series are not always available, and the tSNR may be biased by the presence of (patho-)physiological signal changes and head motion. Therefore, the option implemented in ExploreASL is to use sequence-specific template masks obtained from previous population analyses, after which individual masks are restricted to (pGM+pWM)>0.5 to remove voxels outside the brain. Further development is needed to create masks that take individual anatomical differences in skull and air cavities into account. Noteworthy, ExploreASL applies this analysis mask only for analyses, not for visual QC.

### 4. Population module

This module performs group-level QC and creates group-level results for statistical analyses. Whereas the above-described Structural and ASL modules perform image processing on the individual level, this module performs its analysis on multiple-subjects and/or multi time-point level. For this purpose, ASL images are transformed into standard space using the T1w transformation fields smoothed to the effective spatial resolution of ASL. For transformation of all intermediate and final images, all previous spatial transformations are merged into a single combined transformation to minimize accumulation of interpolation artifacts through the pipeline. Partial volumes of GM and WM obtained from anatomical images are multiplied by the Jacobian determinants of the deformation fields - a.k.a. modulation - to account for voxel-volume changes when transforming to standard space (Ashburner and Friston 1999).

#### 4.1. Template creation

Population templates can reveal population- or sequence-specific perfusion patterns that are not visible on the individual level. ExploreASL generates between-subject mean, standard deviation (SD), CoV, and SNR images for the total study population and, optionally, for different sets (e.g. different centers/sequences/cohorts) within the study (Figure 2). In addition to CBF itself, auxiliary images (e.g. M0), intermediate images (e.g. mean control images), or QC images (e.g. temporal SD) can provide a valuable overview of the data, for example when comparing data originating from different centers.

**Figure 2.**
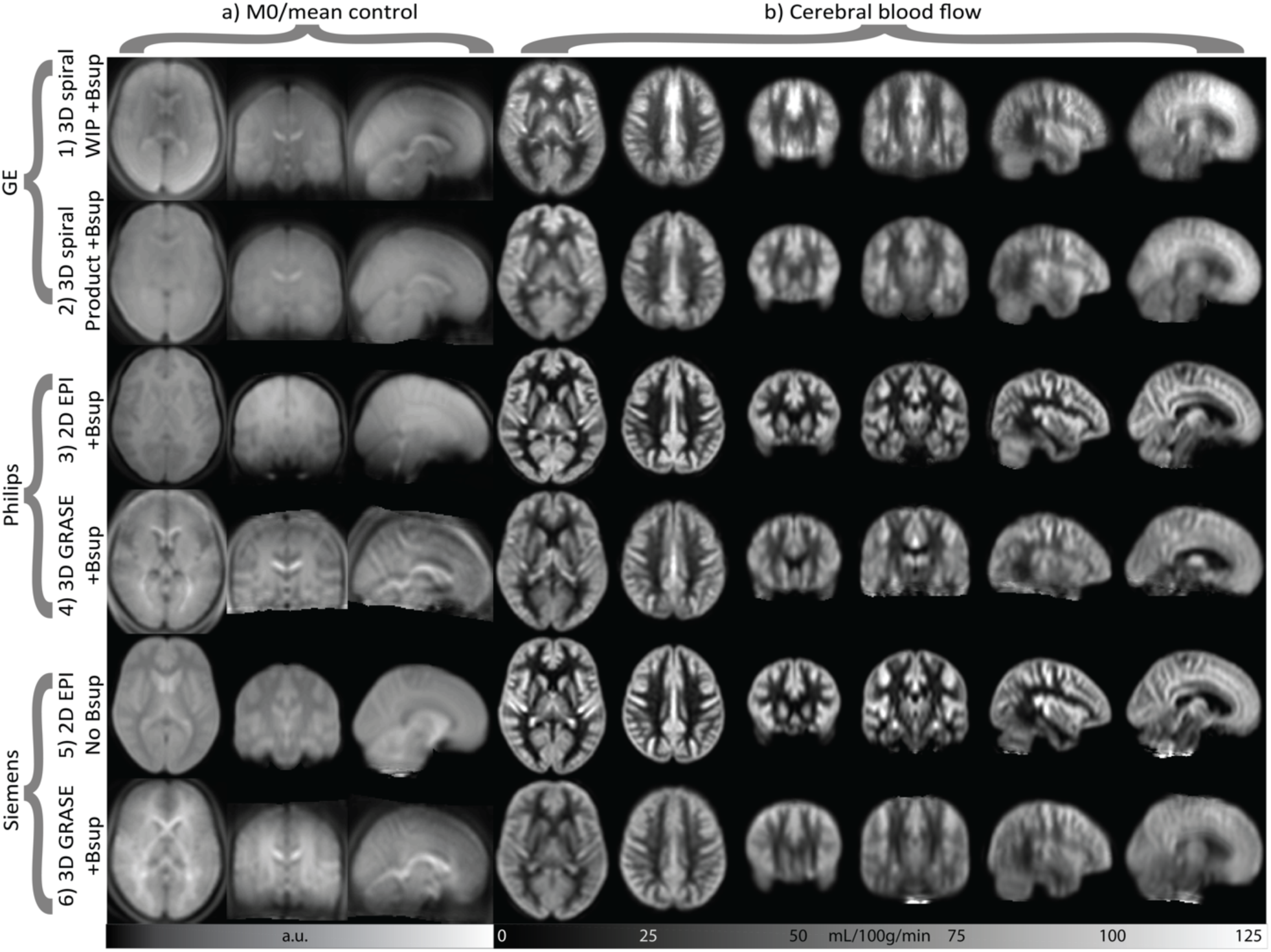
Templates (population-averages from previous studies) are shown for source images (a) and CBF maps (b), for several arterial spin labeling (ASL) acquisitions with/without background suppression (Bsup) from different vendors. The average CBF images are intensity normalized to a mean total GM CBF of 60 mL/min/100g (see Suppl Fig 10 for the unscaled CBF images). Source images are mean control images for Philips and Siemens and M0 images for GE, which does not output control images. Note that the images differ mostly in their effective spatial resolution, orbitofrontal signal dropout, and the amount of macro-vascular artifacts. The differences in geometric distortion are mostly too subtle to be noted on these population-averages images. Note the inferior-superior gradient in the source images in the 2D EPI sequence with background suppression. a.u. = arbitrary units, Bsup = background suppression, WIP = work-in-progress pre-release version. See sequence details in Supplementary Table 1.

#### 4.2. Multi-sequence equalization

Quantitative CBF images can differ between centers because of a number of hardware, labeling, and readout choices implemented by different MRI vendors and/or laboratories (Deibler et al. 2008; Heijtel et al. 2014; Alsop et al. 2015; Jack et al. 2010). Some of these differences can be accounted for, as detailed in the previous sections. However, until a more robust procedure is devised - e.g. the use of a flow phantom (Oliver-Taylor et al. 2017) - a pragmatic approach is required to remove the remaining CBF quantification differences between sequences, scanner types, and centers (Mutsaerts et al. 2019). ExploreASL optionally performs a spatially varying intensity normalization by computing a smooth average CBF bias field for each ASL sequence (Supplementary Figure 10). This step assumes that the studied physiological effects are equally distributed across the subjects scanned with each sequence, scanner and/or site. The final CBF images in the standard space are smoothed with an 8×8×8 mm full-width at half-maximum Gaussian, and averaged to create a sequence/scanner type/site-specific mean CBF image. These sequence-mean CBF images are intensity normalized to GM CBF of 60 mL/100g/min and averaged to create a general mean CBF image. The sequence bias field is calculated by dividing the general mean with the sequence-mean CBF image. The individual CBF images are multiplied by their sequence bias to normalize the intensities across sequences (Mutsaerts et al. 2018).

#### 4.3. ROI statistics

In ExploreASL, ROI masks are created by combining existing atlases with individual GM and WM masks. The GM atlases currently implemented are: (i) MNI structural (Mazziotta et al. 2001), (ii) Harvard-Oxford (Desikan et al. 2006), and (iii) Hammers (Hammers et al. 2002). A deep WM atlas is created by eroding the SPM12 WM tissue class by a 4 voxel sphere (i.e. 6 mm), to avoid signal contamination from the GM (Mutsaerts et al. 2014). Other existing, or custom, atlases can be easily applied. The Online Brain Atlas Reconciliation Tool (OBART) at obart.brainarchitecture.org (Bohland et al. 2009) provides an overview of the overlap and differences between atlases. For each ROI, statistics are calculated separately within the left and right hemisphere, as well as for the full ROI; both with and without PVC. The same CBF statistics are also calculated for user-provided ROIs and lesion masks, as well as for the 25 mm margin around the ROI/lesion (Moghaddasi et al. 2015), for the ipsilateral hemisphere excluding the lesion, and for the same three masks at the contralateral side. Subject-specific ROI and lesion masks are treated the same, except for the fact that lesion masks are also used for the cost function masking (see section 1.3). Individual vascular masks are used to exclude regions with intra-vascular signal (see section 3.7) from CBF statistics, but not from spatial CoV statistics.

Finally, all masks are intersected with a group-level analysis mask, created from the individual analysis masks created in section 3.7. Individual differences of these analysis masks can be caused by differences in head position, FoV, and nasal sinus size. To limit the effects of this mask heterogeneity on statistical analyses, ExploreASL creates a group-level analysis mask from standard-space voxels present in at least 95% of the individuals masks (Supplementary Figure 8).

#### 4.4. Quality Control

On a participant level, ExploreASL outputs QC parameters in a JSON file and provides unmasked images in standard space for visual QC, for both intermediate and final images (Supplementary Figure 11) to detect technical failure, outliers and artifacts. QC parameters are also obtained by comparing individual ASL images with an atlas, a group average, or an average from a previous study. Whole-brain and regional differences larger than 2-3 SD are indicated and should be visually inspected. Deviations can hint to software updates or different scanners and, if not accounted for, can lead to low power of the statistical analyses (Chenevert et al. 2014). All QC parameters and images are also collected in a PDF file (Figure 3, Supplementary Table 2). While these QC parameters can be helpful in detecting artifacts and/or protocol deviations, their use has not yet been validated, and the normal and abnormal range for each of the parameters still need to be determined.

**Figure 3.**
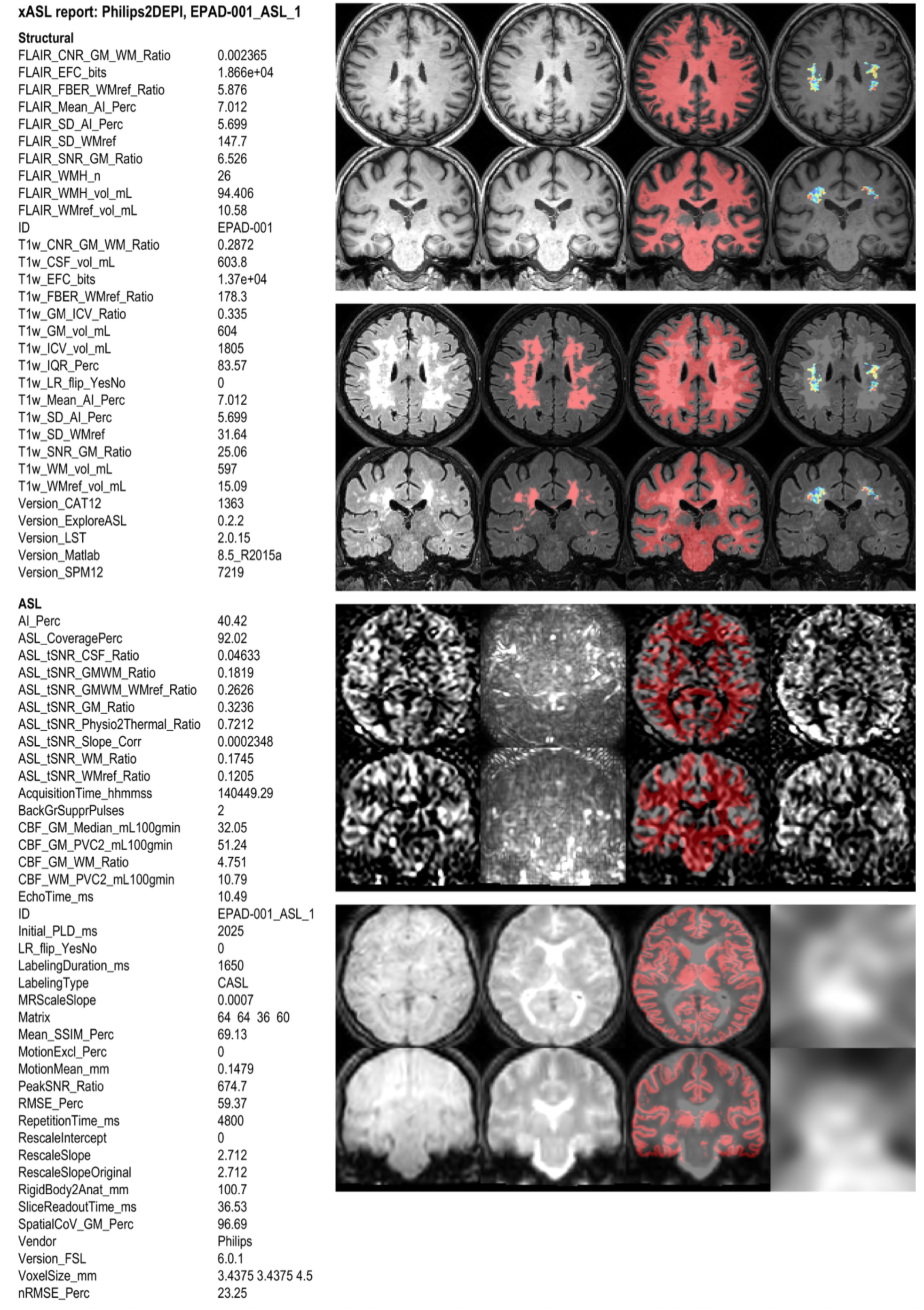
Example PDF report for a single subject. This provenance and QC report includes information collected from each image processing step across the pipeline and assembled in the population module. It is stored in a key-<VALUE> format, facilitating inclusion of plugin or new parameters. Keys and values are grouped into the structural and ASL modules, and the software versions (see Supplementary Table 2). Figures represent transversal and coronal slices in MNI standard space: 1-4) T1w before and after lesion filling, pWM projected over T1w, WMref projected over T1w, 5-8) FLAIR, WMH mask projected over FLAIR, pWM projected over FLAIR, WMref projected over FLAIR, 9-12) CBF, temporal SD, pWM projected over CBF, temporal SNR, 13-16) mean control, M0 before processing, pGM projected over M0, M0 after processing. The pWM/pGM projections in the third column allow a visual assessment of registration performance. CBF = cerebral blood flow, FLAIR = FLuid Attenuated Inversion Recovery, GM = gray matter, pGM = GM partial volume, pWM = WM partial volume, SNR = signal-to-noise ratio, WMref = WM noise reference region, WM = white matter. Example data are from the EPAD study (Ritchie et al. 2016).

## Methods

We illustrate the ExploreASL image processing results and reproducibility for three populations with similar 2D-EPI PCASL protocols: perinatally-infected HIV children, healthy adults, and elderly with mild cognitive complaints, from the NOVICE (Blokhuis et al. 2017), the Sleep (Elvsåshagen et al. 2018), and the European Prevention of Alzheimer’s Dementia (EPAD) studies (Ritchie et al. 2016), respectively (Supplementary Table 3). All three studies adhered to the Declaration of Helsinki and were approved by the local ethics committees (Academic Medical Center (AMC) in Amsterdam, Norwegian South East Regional Ethics Committee, and VU Medical Center Amsterdam and University of Edinburgh, respectively). Written informed consent was obtained from all participants (or parents of children younger than 12 years for NOVICE). Each participant of the Sleep study received NOK 500 for participation.

The performance of image processing should be comparable between different centers, independent of used hardware and OSes, to allow data pooling and comparison between studies. Here, we investigated the between-center reproducibility of the intermediate and final pipeline results without and with the ExploreASL-specific modifications of the SPM12, CAT12, and LST source code (modifications described below). To this end, a single participant from each study was analyzed: with the lowest GM volume from NOVICE and EPAD (GM/ICV ratio 0.41 and 0.33, respectively), and the highest GM volume (GM/ICV ratio 0.55) from the Sleep study. These three datasets were processed at two centers with the following combinations of OS and MATLAB version: Linux-2018b (HZDR, Dresden, Germany; Linux server, 2.1 GHz Intel Xeon 6130, Ubuntu 5), Windows-2015a and 2018b (Amsterdam UMC, The Netherlands; Dell Alienware laptop, 2.9-4.3 GHz Intel i7-7820HK, Windows 10 Version 1903). After each pipeline step, the between-center - or between-system - reproducibility was obtained as difference of the image intensities and orientation between the NIfTIs of the two compared systems. The image intensity reproducibility was calculated as the median voxel-wise relative intensity difference (Kurth et al. 2015), whereas the image orientation reproducibility was calculated as the mean voxel-wise net displacement vector in real-world coordinates (Power et al. 2012). These were calculated for T1w with GM segmentation, FLAIR with WMH segmentation, M0, quantified CBF, GM partial volume in ASL native space (pGM_ASL_), and PV-corrected GM CBF.

More complex calculations involving floating-point arithmetic operations, e.g. matrix inversions, can produce different results between OSes and MATLAB versions in the last digits. These minimal differences can accumulate in iterative algorithms such as segmentation and registration, and propagate across the pipeline. To mitigate these effects, during the process of implementing and using the pipeline for previous clinical studies, we modified parts of the SPM12, CAT12, and LST toolboxes: e.g. using the MATLAB ‘\’ operator for solving a system of linear equations instead of calculating a matrix inversion, providing a separate C++ implementation for convolutions, and/or rounding some calculations to 15 significant digits.

## Results

Running time for a single EPAD participant took 22:11, 4:34, and 0:40 min for the Structural, ASL, and Population modules, respectively (27:25 min in total). On low quality, the same processing took 7:30 min, 2:24, and 0:34 respectively (10:28 min in total) (Windows-2018b). Figure 4 shows differences between populations or sequences on the ExploreASL population-specific parametric maps. While the GM CBF was highest in the pediatric and lowest in the geriatric population (Figure 4c), both the between-subject CoV and within-scan temporal SD were comparable in these populations and lowest in the healthy adults (Figure 4d-e). The temporal SD (Figure 4e) was high in vascular regions and highest around the ventricles in the pediatric dataset, due to a 2D EPI fat-saturation related artefact. Despite these differences, the temporal SNR appeared relatively comparable (Figure 4f), albeit slightly higher for the pediatric population. The average mean control images (Figure 4g) showed subtle differences in background suppression efficacy, as different tissue contrast and inferior-superior background suppression efficiency gradient. Only slight differences in ventricle and sulci size were visible between the pediatric and geriatric population (Figure 4a) confirming satisfactory performance of spatial normalization.

**Figure 4.**
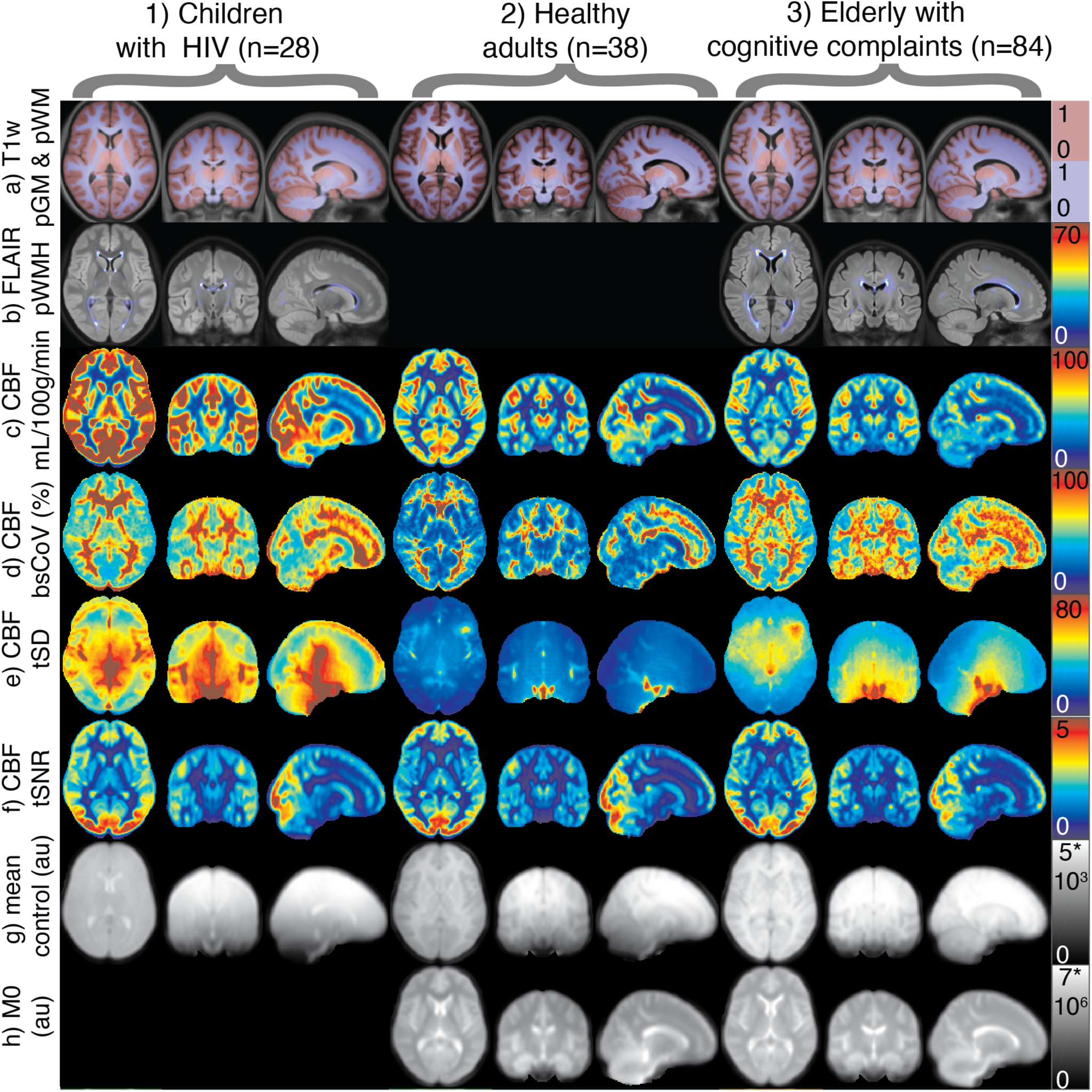
Transversal, coronal, and sagittal population average images for the three example populations: 1) NOVICE, 2) Sleep study, 3) EPAD (see Supplementary Table 3): a) T1w anatomical image with pGM and pWM overlay, b) FLAIR anatomical image overlaid with probability of WMH presence across the whole population, c) Mean CBF, d) between-subject CBF variation (SD CBF/mean CBF per voxel across all subjects), e) temporal SD of CBF, mean over all subjects is shown, f) temporal SNR of CBF (mean CBF/tSD CBF), g) mean control image (note the background suppression gradient), h) M0 calibration image. Note that the FLAIR and M0 were not acquired in the Sleep and NOVICE studies, respectively. CBF = cerebral blood flow, tSNR = temporal signal-to-noise ratio, tSD = temporal standard deviation, au = arbitrary units, bs = between-subject, CoV = coefficient of variance, p = probability, GM = gray matter, WM = white matter, WMH = white matter hyperintensity.

All three datasets showed zero difference when the pipeline was repeated twice on the same system. When comparing OSes only - Linux-2018b vs Windows-2018b - the structural module showed final voxel-wise differences of 0.77% pGM in NOVICE and 1.74% WMH in EPAD that became negligible after our code modifications (Figure 5). The ASL module differences were smaller than 0.5%, except for the pGM_ASL_ (0.57-2.5%) and PV-corrected GM CBF (0.61-5.76%). Both improved after modifications to 0-1.2% and 0.32-1.5% for pGM_ASL_ and PV-corrected GM CBF, respectively, showing the impact of our modifications. The reproducibility between OS and MATLAB versions - Linux-2018b vs Windows-2015a - showed satisfactory post-modification reproducibility, e.g. pGM_ASL_ (0.47-1.79%) and GM CBF (0.57-1.77%) (Supplementary Table 4). Compared with the above-mentioned Linux-2018b vs Windows 2018b results this shows an additional decrease in reproducibility when a different MATLAB version is used on top of different OSes and/or systems.

**Figure 5.**
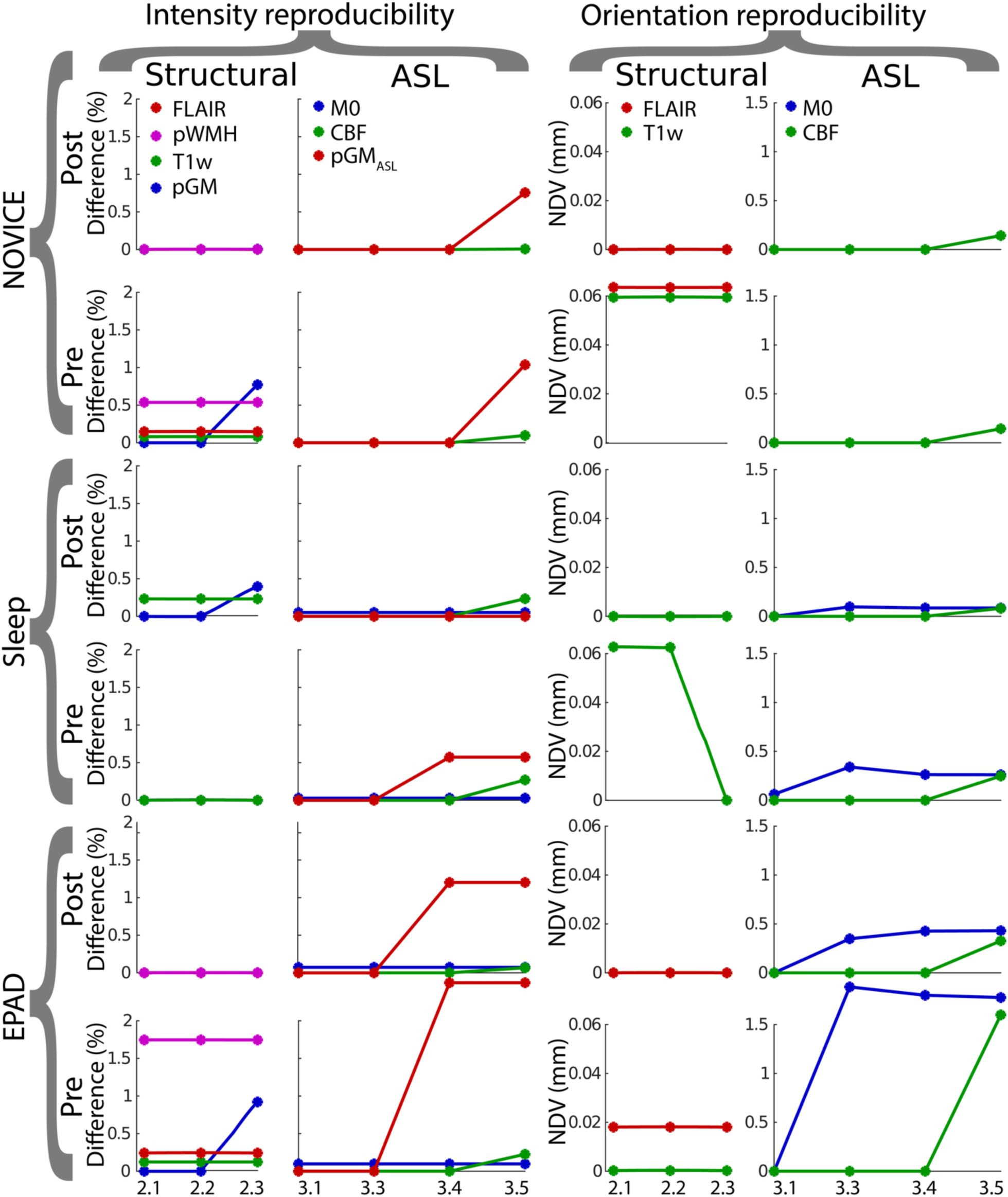
Reproducibility between Linux-2018b and Windows-2018b and for the three datasets NOVICE, Sleep, and EPAD. Results are shown before (pre) and after (post) the ExploreASL-specific modifications of MATLAB and SPM12 code. The median relative intensity difference is shown in the two columns on the left (referred to as difference) and the mean voxel-wise net displacement vector (NDV) is shown in the two columns on the right. Labels on the x-axis describe the processing steps in the Structural (2.1 = WMH correction, 2.2 = Segmentation, 2.3 = Spatial normalization) and ASL module (3.1 = Motion correction, 3.3 = T1w-ASL registration, 3.4 = M0 processing, 3.5 = CBF quantification) as described in Table 1. pGMASL = gray matter partial volume map in ASL space.

## Discussion and future directions

In this manuscript, we reviewed many of the most salient ASL image processing choices, and their implementation in ExploreASL version 1.0.0. We demonstrated the software’s functionality to review individual cases as well as population-average images for quality control. Our findings show that between-center computing differences can lead to voxel-wise CBF quantification differences of up to 5.7% on average for the total GM, which were reduced to 1.7% by addressing implementation differences of complex floating-point operations between MATLAB versions and OSes. This may especially be beneficial for multi-center studies or for pooling multiple ASL studies to attain sample sizes required for the discovery of subtle (patho-)physiological perfusion patterns.

Several other ASL image processing pipelines are publicly available and free for academic use, each providing specific features. The first publicly available pipeline ASLtbx quantifies CBF of various ASL sequences (Wang et al. 2008) and features customized motion correction and advanced outlier detection (Dolui et al. 2017); ASAP contains a graphical user interface (GUI) with an interface for population analyses, and generates statistical reports (Mato Abad et al. 2016); the ASLM toolbox is a MATLAB- and SPM-based command-line tool (Homan et al. 2012), ASL-MRICloud features a web interface with an automated cloud solution (Li et al. 2018); ASL-QC handles multiple vendors and provides QC metrics (not published); BASIL uses a Bayesian approach for the quantification and PVC of multi-TI, QUASAR, and time-encoded ASL data, thus offering the most comprehensive quantification (Chappell et al. 2010); CBFBIRN offers an online data repository with online image processing (Shin et al. 2016); Functional ASL (FASL) (FASL webpage) and fMRI Grocer (Zhu Grocer) are SPM toolboxes that process both functional ASL and BOLD MRI; GIN fMRI performs separate control and label realignment and automatically excludes outliers and volumes with strong motion (unpublished); MilxASL features spatial and temporal denoising (Fazlollahi et al. 2015); MJD-ASL is implemented into ‘cranial cloud’, addresses noise concerns and processes cerebral blood volume (CBV) (MJD-ASL Webpage); NiftyFit supports quantification of other MRI sequences as IVIM, NODDI, and relaxometry (Melbourne et al. 2016); VANDPIRE is Python-based, has a scanner console plugin and allows flow territory mapping from vessel-encoded ASL (Arteaga et al. 2017).

The main advantage of ExploreASL is the use of image processing that is optimized for clinical studies, to address the diversity of the clinical populations, hardware, and sequences used. Other strengths include its between-center reproducibility of image processing, and its flexible open source development through GitHub with a growing team of international scientists. This improves the likelihood of rapid debugging, encourages inter-institutional cross-pollination in validation of new techniques, and allows quick adaptation of the software to new regulations regarding e.g. best practice (Nichols et al. 2017) and data transfer (Regulation 2016). Moreover, the use of a common validated pipeline rather than in-house software increases reliability and reproducibility in neuroscience research (Poldrack et al. 2017). The ExploreASL team actively focuses on keeping the pipeline user-friendly by allowing download on demand during the development phase and by being in close-contact with all the users. ExploreASL was built upon freely available software that performs well in a wide array of cases, rather than opting for solutions with optimal performance in specific cases but not applicable in general. Still, the modular structure of ExploreASL allows the replacement of some steps by solutions tailored for specific datasets.

Although ExploreASL allows custom labeling efficiency values and global CBF calibration, it does not estimate labeling efficiency. While literature values for labeling efficiency (Dai et al. 2008) may suffice in many clinical cases (Heijtel et al. 2014), individual correction can be beneficial for specific populations (Václavů et al. 2018). For these, phase-contrast MRI can improve the CBF quantification by a) calibrating CBF based on total flow through the brain-feeding arteries (Aslan and Lu 2010; Ambarki et al. 2015), or b) modeling the labeling efficiency based on the velocity in the labeling plane (Václavů et al. 2019). Compared to ASL, drawbacks of the phase-contrast MRI include its lower reproducibility for whole brain CBF estimates (Dolui et al. 2016), and its lower agreement with PET (Puig et al. 2018). Moreover, an automatic implementation requires good data quality, perpendicular placement of the labeling plane to the vessels, and the absence of vessel tortuosity, conditions that are rarely met in clinical datasets. Future solutions may be provided by new sequences under development, which allow direct labeling efficiency measurements during the ASL acquisition (Chen et al. 2018; Lorenz et al. 2018).

Several additional features are scheduled for future releases, after being tested first through the GitHub beta versions of ExploreASL. These include full BIDS support (Gorgolewski et al. 2016); support for Hitachi and Canon datasets; unit testing to ensure stability of the pipeline through the continuous development; inclusion of WM atlases for extended WM analysis; a GUI for easier configuration and execution; quantification of advanced ASL schemes such as velocity- and acceleration-selective ASL (Schmid et al. 2015) and integration of the BASIL toolbox to allow multi-PLD and time-encoded sequence quantification (Chappell et al. 2009); and support for individual-center calibration, e.g. using the recently introduced Quantitative ASL Perfusion Reference (QASPER) (Oliver-Taylor et al. 2017) phantom (Gold Standard Phantoms, London, UK). Although ExploreASL’s computation times are moderate for research purposes, a clinical scanner implementation would benefit from parallelization on graphical processing units (GPUs) to provide robust automatic QC within clinical scanning time (e.g. <5 min). Another improvement would be the investigation of the effect of image processing choices, as well as the availability of physiological and quantification parameters for different populations (Fazlollahi et al. 2015). This would allow for the incorporation of quantification confidence intervals in the output of ExploreASL. For anonymization purposes, the face can be removed from the structural scans (Nichols et al. 2017; Leung et al. 2015) using a defacing algorithm such as the one implemented in SPM12, but further testing is required to verify that the analysis is not affected (de Sitter et al. 2017). Statistical analyses can be biased for populations with large inter-subject differences in their deformations, e.g., developing brains or a wide range of atrophy. The CerebroMatic toolbox (Wilke et al. 2017) is a tool that accounts for this bias and will be incorporated in future releases of ExploreASL. Finally, we intend to implement ExploreASL as a cloud solution and scanner console plugin.

Image processing techniques that require validation include: using the UNWARP toolbox for simultaneous motion and susceptibility deformation correction (Andersson et al. 2001), using temporal information for artifact removal through the application of an independent-component analysis (ICA) (Wells et al. 2010; Hao et al. 2018; Zhu et al. 2018) or using respiratory and cardiac signal (Restom et al. 2006), spatial denoising - once validated under realistic conditions (Spann et al. 2017; Wells et al. 2010; Bibic et al. 2010; Liang et al. 2015), obtaining GM-WM segmentations from fractional signal modeling of the magnetization recovery profile acquired with a Look-Locker readout (Petr et al. 2013; Ahlgren et al. 2014), and using the BBR method for motion correction or registration (Greve and Fischl 2009). Finally, we aim to improve the inter-center reproducibility even further.

### Conclusion

ExploreASL is a versatile pipeline that performs well on a wide-range of diseases, including datasets with lesions, allows flexible parameter definition, and a quick exploration of datasets and QC images of each pipeline step in the same space. We made the pipeline available at www.ExploreASL.com. We anticipate that ExploreASL will allow for more flexible collaboration amongst clinicians and scientists, help to achieve the consensus standards for ASL processing sought by the OSIPI, facilitate validation of ASL image processing approaches, and accelerate translation to clinical research and practice.

## Acknowledgments

This project has received support from the following EU/EFPIA Innovative Medicines Initiatives (1 and 2) Joint Undertakings: EPAD grant no. 115736, AMYPAD grant no. 115952. Additionally, this work received support from the EU-EFPIA Innovative Medicines Initiatives Joint Undertaking (grant No 115952). HM is supported by Amsterdam Neuroscience funding. FB and XG are supported by NIHR funding through the UCLH Biomedical Research Centre. DLT is supported by the UCL Leonard Wolfson Experimental Neurology Centre (PR/ylr/18575). EDV is supported by the Wellcome/EPSRC Centre for Medical Engineering [WT 203148/Z/16/Z]. IA is supported by The Gleason Foundation. MJPvO receives research support from Philips, the EU under the Horizon 2020 program (project: CDS-QUAMRI, project number 634541), and the research program Innovational Research Incentives Scheme Vici with project number 016.160.351, which is financed by the Netherlands Organization for Scientific Research (NWO). MC received funding from the Engineering and Physical Sciences Research Council UK (EP/P012361/1). The Wellcome Centre for Integrative Neuroimaging is supported by core funding from the Wellcome Trust (203139/Z/16/Z). The authors wish to thank the COST-AID (European Cooperation in Science and Technology - Arterial spin labeling Initiative in Dementia) Action BM1103 and the Open Source Initiative for Perfusion Imaging (OSIPI) and the ISMRM Perfusion Study groups for facilitating meetings for researchers to discuss the implementation of ExploreASL. The authors acknowledge Guillaume Flandin, Robert Dahnke, and Paul Schmidt for reviewing the structural module for its implementation of SPM12, CAT12, and LST, respectively; Krzysztof Gorgolewksi for his advice on the BIDS implementation; Jens Maus for help with MEX compilation; Cyril Pernet for providing the SPM Univariate Plus scripts; and Koen Baas for curating the Philips 3D GRASE data. The authors acknowledge the following researchers and teams: Yannis Paloyelis from King’s College London, for providing the data of the INtranasal OxyTocin trial, Torbjørn Elvsåshagen from Oslo University Hospital for providing the Sleep study dataset; the EPAD investigators for providing the Amsterdam site elderly dataset; Kim van de Ven from Philips Healthcare for providing the 3D GRASE dataset; Philip de Witt Hamer from Amsterdam UMC for providing the PICTURE dataset, and Chris Chen from the National University of Singapore for providing the Singapore Memory Clinical dataset.

## References

Ahlgren, A., Wirestam, R., Knutsson, L., Petersen, E.T., 2018. Improved calculation of the equilibrium magnetization of arterial blood in arterial spin labeling. Magn. Reson. Med. 1–9.

Ahlgren, A., Wirestam, R., Thade, E., Ståhlberg, F., Knutsson, L., Petersen, E.T., Ståhlberg, F., Knutsson, L., 2014. Partial volume correction of brain perfusion estimates using the inherent signal data of time-resolved arterial spin labeling. NMR Biomed. 27, 1112–1122.

Almeida, J.R.C., Greenberg, T., Lu, H., Chase, H.W., Jay, C., Cooper, C.M., Deckersbach, T., Adams, P., Carmody, T., Fava, M., Kurian, B., Mcgrath, P.J., Mcinnis, M.G., Oquendo, M.A., Parsey, R., Weissman, M., Trivedi, M., Phillips, M.L., Hospital, M.G., Arbor, A., Brook, S., 2018. Test-retest reliability of cerebral blood flow in healthy individuals using arterial spin labeling: Findings from the EMBARC study. Magn. Reson. Imaging 45, 26–33.

Alsop, D.C., Detre, J.A., 1999. Background suppressed 3D RARE ASL perfusion imaging, in: International Society for Magnetic Resonance in Medicine. Philadelphia, p. 601.

Alsop, D.C., Detre, J.A., 1996. Reduced transit-time sensitivity in noninvasive magnetic resonance imaging of human cerebral blood flow. J. Cereb. Blood Flow Metab. 16, 1236–1249.

Alsop, D.C., Detre, J.A., Golay, X., Günther, M., Hendrikse, J., Hernandez-Garcia, L., Lu, H., MacIntosh, B.J., Parkes, L.M., Smits, M., van Osch, M.J.P., Wang, D.J.J., Wong, E.C., Zaharchuk, G., 2015. Recommended implementation of arterial spin-labeled perfusion MRI for clinical applications: A consensus of the ISMRM perfusion study group and the European consortium for ASL in dementia. Magn. Reson. Med. 73, 102–116.

Alsop, D.C., Detre, J.A., Grossman, M., 2000. Assessment of cerebral blood flow in Alzheimer’s disease by spin-labeled magnetic resonance imaging. Ann.Neurol. 47, 93–100.

Ambarki, K., Wåhlin, A., Zarrinkoob, L., Wirestam, R., Petr, J., Malm, J., Eklund, A., 2015. Accuracy of Parenchymal Cerebral Blood Flow Measurements Using Pseudocontinuous Arterial Spin-Labeling in Healthy Volunteers. AJNR Am. J. Neuroradiol. 36, 1816–1821.

Andersson, J.L.R., Hutton, C., Ashburner, J., Turner, R., Friston, K., 2001. Modeling geometric deformations in EPI time series. Neuroimage 13, 903–919.

Andersson, J.L.R., Skare, S., Ashburner, J., 2003. How to correct susceptibility distortions in spin-echo echo-planar images: application to diffusion tensor imaging. Neuroimage 20, 870–888.

Arteaga, D.F., Strother, M.K., Davis, L.T., Fusco, M.R., Faraco, C.C., Roach, B.A., Scott, A.O., Donahue, M.J., 2017. Planning-free cerebral blood flow territory mapping in patients with intracranial arterial stenosis. J. Cereb. Blood Flow Metab. 37, 1944–1958.

Ashburner, J., 2012. SPM: A history. Neuroimage 62, 791–800.

Ashburner, J., 2007. A fast diffeomorphic image registration algorithm. Neuroimage 38, 95–113.

Ashburner, J., Friston, K.J., 2011. Diffeomorphic registration using geodesic shooting and Gauss–Newton optimisation. Neuroimage 55, 954–967.

Ashburner, J., Friston, K.J., 2005. Unified segmentation. Neuroimage 26, 839–851.

Ashburner, J., Friston, K.J., 1999. Nonlinear spatial normalization using basis functions. Hum.Brain Mapp.

Ashburner, J., Ridgway, G.R., 2012. Symmetric diffeomorphic modeling of longitudinal structural MRI. Front. Neurosci. 6, 197.

Aslan, S., Lu, H., 2010. On the sensitivity of ASL MRI in detecting regional differences in cerebral blood flow. Magn. Reson. Imaging 28, 928–935.

Asllani, I., Borogovac, A., Brown, T.R., 2008. Regression algorithm correcting for partial volume effects in arterial spin labeling MRI. Magn. Reson. Med. 60, 1362–1371.

Asllani, I., Habeck, C., Borogovac, A., Brown, T.R., Brickman, A.M., Stern, Y., 2009. Separating function from structure in perfusion imaging of the aging brain. HBM 30, 2927–2935.

Battaglini, M., Jenkinson, M., De Stefano, N., 2012. Evaluating and reducing the impact of white matter lesions on brain volume measurements. Hum. Brain Mapp. 33, 2062–2071.

Beaumont, H., 2015. Multimodal Magnetic Resonance Imaging of Frontotemporal Lobar Degeneration.

Bibic, A., Knutsson, L., Ståhlberg, F., Wirestam, R., 2010. Denoising of arterial spin labeling data: wavelet-domain filtering compared with Gaussian smoothing. MAGMA 23, 125–137.

Blokhuis, C., Mutsaerts, H.J.M.M., Cohen, S., Scherpbier, H.J., Caan, M.W.A., Majoie, C.B.L.M., Kuijpers, T.W., Reiss, P., Wit, F.W.N.M., Pajkrt, D., 2017. Higher subcortical and white matter cerebral blood flow in perinatally HIV-infected children. Medicine 96, e5891.

Bohland, J.W., Bokil, H., Allen, C.B., Mitra, P.P., 2009. The Brain Atlas Concordance Problem: Quantitative Comparison of Anatomical Parcellations. PLoS One 4, e7200.

Bron, E.E., Steketee, R.M.E., Houston, G.C., Oliver, R.A., Achterberg, H.C., Loog, M., van Swieten, J.C., Hammers, A., Niessen, W.J., Smits, M., Klein, S., Alzheimer’s Disease Neuroimaging Initiative, 2014. Diagnostic classification of arterial spin labeling and structural MRI in presenile early stage dementia. Hum. Brain Mapp. 35, 4916–4931.

Cardoso, M.J., Modat, M., Vercauteren, T., Ourselin, S., 2015. Scale Factor Point Spread Function Matching: Beyond Aliasing in Image Resampling, in: Medical Image Computing and Computer-Assisted Intervention -- MICCAI 2015. Springer International Publishing, pp. 675–683.

Çavuşoğlu, M., Pfeuffer, J., Uğurbil, K., Uludağ, K., 2009. Comparison of pulsed arterial spin labeling encoding schemes and absolute perfusion quantification. Magn. Reson. Imaging 27, 1039–1045.

Chappell, M.A., Groves, A.R., Whitcher, B., Woolrich, M.W., 2009. Variational Bayesian Inference for a Nonlinear Forward Model. IEEE Trans. Signal Process. 57, 223–236.

Chappell, M.A., MacIntosh, B.J., Donahue, M.J., Günther, M., Jezzard, P., Woolrich, M.W., 2010. Separation of macrovascular signal in multi-inversion time arterial spin labelling MRI. Magn. Reson. Med. 63, 1357–1365.

Chappell, M., MacIntosh, B., Okell, T., 2018. Introduction to Perfusion Quantification using Arterial Spin Labelling. Oxford University Press.

Chard, D.T., Jackson, J.S., Miller, D.H., Wheeler-Kingshott, C.A.M., 2010. Reducing the impact of white matter lesions on automated measures of brain gray and white matter volumes. J. Magn. Reson. Imaging 32, 223– 228.

Chenevert, T.L., Malyarenko, D.I., Newitt, D., Li, X., Jayatilake, M., Tudorica, A., Fedorov, A., Kikinis, R., Liu, T.T., Muzi, M., Oborski, M.J., Laymon, C.M., Li, X., Thomas, Y., Jayashree, K.-C., Mountz, J.M., Kinahan, P.E., Rubin, D.L., Fennessy, F., Huang, W., Hylton, N., Ross, B.D., 2014. Errors in Quantitative Image Analysis due to Platform-Dependent Image Scaling. Transl. Oncol. 7, 65–71.

Chen, J.J., Rosas, H.D., Salat, D.H., 2011. Age-associated reductions in cerebral blood flow are independent from regional atrophy. Neuroimage 55, 468–478.

Chen, Y., Wang, D.J.J., Detre, J.A., 2011. Test-retest reliability of arterial spin labeling with common labeling strategies. J. Magn. Reson. Imaging 33, 940–949.

Chen, Z., Zhao, X., Zhang, X., Guo, R., Teeuwisse, W.M., Zhang, B., Koken, P., Smink, J., Yuan, C., van Osch, M.J.P., 2018. Simultaneous measurement of brain perfusion and labeling efficiency in a single pseudo-continuous arterial spin labeling scan. Magn. Reson. Med. 79, 1922–1930.

Clement, P., Mutsaerts, H., Vaclavu, L., Ghariq, E., Pizzini, F.B., Smits, M., Acou, M., Jovicich, J., Vanninen, R., Kononen, M., Wiest, R., Rostrup, E., Bastos-Leite, A.J., Larsson, E.-M.M., Achten, E., 2017. Variability of physiological brain perfusion in healthy subjects - A systematic review of modifiers. Considerations for multi-center ASL studies. J. Cereb. Blood Flow Metab. In press. https://doi.org/10.1177/0271678X17702156

Crinion, J., Ashburner, J., Leff, A., Brett, M., Price, C., Friston, K., 2007. Spatial normalization of lesioned brains: Performance evaluation and impact on fMRI analyses. Neuroimage 37, 866–875.

Croal, P.L., Kennedy-McConnel, F., Harris, B., Ma, R., Ng, S.M., Plaha, P., Lord, S., Sibson, N.R., Chappel, M.A., 2019. Quantification of Cerebral Blood Flow using arterial spin labeling in glioblastoma multiforme; challenges of calibration in the presence of oedema, in: Proceedings of ISMRM 2019. Presented at the International Society for Magnetic Resonance in Medicine, p. 2315.

Dai, W., Garcia, D., De, B.C., Alsop, D.C., 2008. Continuous flow-driven inversion for arterial spin labeling using pulsed radio frequency and gradient fields. Magn Reson.Med. 60, 1488–1497.

Deibler, A.R., Pollock, J.M., Kraft, R.A., Tan, H., Burdette, J.H., Maldjian, J.A., 2008. Arterial spin-labeling in routine clinical practice, part 1: technique and artifacts. AJNR Am. J. Neuroradiol. 29, 1228–1234.

Desikan, R.S., Ségonne, F., Fischl, B., Quinn, B.T., Dickerson, B.C., Blacker, D., Buckner, R.L., Dale, A.M., Maguire, R.P., Hyman, B.T., Albert, M.S., Killiany, R.J., 2006. An automated labeling system for subdividing the human cerebral cortex on MRI scans into gyral based regions of interest. Neuroimage 31, 968–980.

de Sitter, A., Steenwijk, M.D., Ruet, A., Versteeg, A., Liu, Y., van Schijndel, R.A., Pouwels, P.J.W., Kilsdonk, I.D., Cover, K.S., van Dijk, B.W., Ropele, S., Rocca, M.A., Yiannakas, M., Wattjes, M.P., Damangir, S., Frisoni, G.B., Sastre-Garriga, J., Rovira, A., Enzinger, C., Filippi, M., Frederiksen, J., Ciccarelli, O., Kappos, L., Barkhof, F., Vrenken, H., 2017. Performance of five research-domain automated WM lesion segmentation methods in a multi-center MS study. Neuroimage 163, 106–114.

de Sitter, A., Visser, M., Brouwer, I., van Schijndel, R.A., Uitdehaag, B.M.J., Barkhof, F., Vrenken, H., 2017. Impact of removing facial features from MR images of MS patients on automatic lesion and atrophy metrics. Multiple Sclerosis Journal 23, 226–226.

Detre, J.A., Alsop, D.C., Vives, L.R., Maccotta, L., Teener, J.W., Raps, E.C., 1998. Noninvasive MRI evaluation of cerebral blood flow in cerebrovascular disease. Neurology 50, 633–641.

Detre, J.A., Leigh, J.S., Williams, D.S., Koretsky, A.P., 1992. Perfusion imaging. Magn. Reson. Med. 23, 37–45.

De Vis, J.B., Hendrikse, J., Groenendaal, F., de Vries, L.S., Kersbergen, K.J., Benders, M.J.N.L., Petersen, E.T., 2014. Impact of neonate haematocrit variability on the longitudinal relaxation time of blood: Implications for arterial spin labelling MRI. NeuroImage: Clinical 4, 517–525.

Dolui, S., Wang, Z., Shinohara, R.T., Wolk, D.A., Detre, J.A., Alzheimer’s Disease Neuroimaging Initiative, 2017. Structural Correlation-based Outlier Rejection (SCORE) algorithm for arterial spin labeling time series. J. Magn. Reson. Imaging 45, 1786–1797.

Dolui, S., Wang, Z., Wang, D.J.J., Mattay, R., Finkel, M., Elliott, M., Desiderio, L., Inglis, B., Mueller, B., Stafford, R.B., Launer, L.J., Jacobs, D.R., Bryan, R.N., Detre, J.A., 2016. Comparison of non-invasive MRI measurements of cerebral blood flow in a large multisite cohort. J. Cereb. Blood Flow Metab. 36, 1244–1256.

Elvsåshagen, T., Mutsaerts, H.J., Zak, N., Norbom, L.B., Quraishi, S.H., Pedersen, P.Ø., Malt, U.F., Westlye, L.T., Van Someren, E.J., Bjørnerud, A., Groote, I.R., 2018. Cerebral blood flow changes after a day of wake, sleep, and sleep deprivation. Neuroimage. https://doi.org/S1053811918321104

Evans, A.C., Janke, A.L., Collins, D.L., Baillet, S., 2012. Brain templates and atlases. Neuroimage 62, 911–922.

Fallis, A.G., 2013. Best Practices in Data Analysis and Sharing in Neuroimaging using MRI. J. Chem. Inf. Model. 53, 1689–1699.

FASL Webpage, http://web.eecs.umich.edu/~hernan/Public/Programs/ (accessed 6.18.19).

Fazlollahi, A., Bourgeat, P., Liang, X., Meriaudeau, F., Connelly, A., Salvado, O., Calamante, F., 2015. Reproducibility of multiphase pseudo-continuous arterial spin labeling and the effect of post-processing analysis methods. Neuroimage 117, 191–201.

Flandin, G., Friston, K., 2008. Statistical parametric mapping (SPM). Scholarpedia J. 3, 6232.

Gai, N.D., Yu, Y., Pham, D., Butman, J.A., 2017. Reduced distortion artifact whole brain CBF mapping using blip-reversed non-segmented 3D echo planar imaging with pseudo-continuous arterial spin labeling. Magn. Reson. Imaging 44, 119–124.

Gaser, C., 2009. Partial Volume Segmentation with Adaptive Maximum A Posteriori (MAP) Approach. Neuroimage 47, S121.

Gevers, S., Van Osch, M.J., Bokkers, R.P.H., Kies, D.A., Teeuwisse, W.M., Majoie, C.B., Hendrikse, J., Nederveen, A.J., 2011. Intra-and multicenter reproducibility of pulsed, continuous and pseudo-continuous arterial spin labeling methods for measuring cerebral perfusion. J. Cereb. Blood Flow Metab. https://doi.org/10.1038/jcbfm.2011.10

Gorgolewski, K.J., Auer, T., Calhoun, V.D., Craddock, R.C., Das, S., Duff, E.P., Flandin, G., Ghosh, S.S., Glatard, T., Halchenko, Y.O., Handwerker, D.A., Hanke, M., Keator, D., Li, X., Michael, Z., Maumet, C., Nichols, B.N., Nichols, T.E., Pellman, J., Poline, J.-B., Rokem, A., Schaefer, G., Sochat, V., Triplett, W., Turner, J.A., Varoquaux, G., Poldrack, R.A., 2016. The brain imaging data structure, a format for organizing and describing outputs of neuroimaging experiments. Scientific Data 3, 160044.

Greve, D.N., Fischl, B., 2009. Accurate and robust brain image alignment using boundary-based registration. Neuroimage 48, 63–72.

Hales, P.W., Kawadler, J.M., Aylett, S.E., Kirkham, F.J., Clark, C.A., 2014. Arterial spin labeling characterization of cerebral perfusion during normal maturation from late childhood into adulthood: normal “reference range” values and their use in clinical studies. J.Cereb.Blood Flow Metab 34, 776–784.

Hales, P.W., Kirkham, F.J., Clark, C.A., 2016. A general model to calculate the spin-lattice (T1) relaxation time of blood, accounting for haematocrit, oxygen saturation and magnetic field strength. J.Cereb.Blood Flow Metab 36, 370–374.

Hammers, A., Koepp, M.J., Free, S.L., Brett, M., Richardson, M.P., Labbé, C., Cunningham, V.J., Brooks, D.J., Duncan, J., 2002. Implementation and application of a brain template for multiple volumes of interest. Hum. Brain Mapp. 15, 165–174.

Handley, R., Zelaya, F.O., Reinders, A.A.T.S., Marques, T.R., Mehta, M.A., O’Gorman, R., Alsop, D.C., Taylor, H., Johnston, A., Williams, S., McGuire, P., Pariante, C.M., Kapur, S., Dazzan, P., 2013. Acute effects of single-dose aripiprazole and haloperidol on resting cerebral blood flow (rCBF) in the human brain. Hum. Brain Mapp. 34, 272–282.

Hao, X., Petr, J., Nederveen, A.J., Wood, J.C., Wang, D.J.J., Mutsaerts, H.J., Jann, K., 2018. ICA cleanup for improved SNR in arterial spin labeling perfusion MRI, in: International Society for Magnetic Resonance in Medicine.

Heijtel, D.F.R., Mutsaerts, H.J.M.M., Bakker, E., Schober, P., Stevens, M.F., Petersen, E.T., van Berckel, B.N.M., Majoie, C.B.L.M., Booij, J., van Osch, M.J.P., van Bavel, E.T., Boellaard, R., Lammertsma, A.A., Nederveen, A.J., 2014. Accuracy and precision of pseudo-continuous arterial spin labeling perfusion during baseline and hypercapnia: a head-to-head comparison with ^15^O H₂O positron emission tomography. Neuroimage 92, 182– 192.

Homan, P., Kindler, J., Hubl, D., Dierks, T., 2012. Auditory verbal hallucinations: imaging, analysis, and intervention. Eur. Arch. Psychiatry Clin. Neurosci. 262 Suppl 2, S91–5.

Jack, C.R., Bernstein, M.A., Borowski, B., Gunter, J.L., Fox, N.C., Thompson, P., Schuff, N., Krueger, G., Killiany, R.J., DeCarli, C., Dale, A.M., Carmichael, O.W., Tosun, D., Weiner, M.W., 2010. Update on the magnetic resonance imaging core of the Alzheimer’s disease neuroimaging initiative. Alzheimers.Dement. 6, 212–220.

Joris, P.J., Mensink, R.P., Adam Tanja C, Liu, T.T., 2018. Cerebral Blood Flow Measurements in Adults: A Review on the Effects of Dietary Factors and Exercise. Nutrients 10, 1–15.

Klein, A., Andersson, J., Ardekani, B.A., Ashburner, J., Avants, B., Chiang, M.-C., Christensen, G.E., Collins, D.L., Gee, J., Hellier, P., Song, J.H., Jenkinson, M., Lepage, C., Rueckert, D., Thompson, P., Vercauteren, T., Woods, R.P., Mann, J.J., Parsey, R.V., 2009. Evaluation of 14 nonlinear deformation algorithms applied to human brain MRI registration. Neuroimage 46, 786–802.

Klein, S., Staring, M., Murphy, K., Viergever, M.A., Pluim, J.P.W., 2010. Elastix: A toolbox for intensity-based medical image registration. IEEE Trans. Med. Imaging 29, 196–205.

Kurth, F., Gaser, C., Luders, E., 2015. A 12-step user guide for analyzing voxel-wise gray matter asymmetries in statistical parametric mapping (SPM). Nat. Protoc. 10, 293–304.

Law, I., Iida, H., Holm, S., Nour, S., Rostrup, E., Svarer, C., Paulson, O.B., 2000. Quantitation of Regional Cerebral Blood Flow Corrected for Partial Volume Effect Using O-15 Water and PET: II. Normal Values and Gray Matter Blood Flow Response to Visual Activation. J. Cereb. Blood Flow Metab. 20, 1252–1263.

Leung, K.Y.E., van der Lijn, F., Vrooman, H.A., Sturkenboom, M.C.J.M., Niessen, W.J., 2015. IT Infrastructure to Support the Secondary Use of Routinely Acquired Clinical Imaging Data for Research. Neuroinformatics 13, 65–81.

Liang, X., Connelly, A., Calamante, F., 2015. Voxel-Wise Functional Connectomics Using Arterial Spin Labeling Functional Magnetic Resonance Imaging: The Role of Denoising. Brain Connect. 5, 543–553.

Liang, X., Connelly, A., Calamante, F., 2013. Improved partial volume correction for single inversion time arterial spin labeling data. Magn. Reson. Med. 69, 531–537.

Liu, H.-L., Kochunov, P., Hou, J., Pu, Y., Mahankali, S., Feng, C.-M., Yee, S.-H., Wan, Y.-L., Fox, P.T., Gao, J.-H., 2001. Perfusion-weighted imaging of interictal hypoperfusion in temporal lobe epilepsy using FAIR-HASTE: Comparison with H215O PET measurements. Magn. Reson. Med. 45, 431–435.

Li, W., Liu, P., Lu, H., Strouse, J.J., van Zijl, P.C.M., Qin, Q., 2017. Fast measurement of blood T 1 in the human carotid artery at 3T: Accuracy, precision, and reproducibility. Magn. Reson. Med. 77, 2296–2302.

Li, X., Morgan, P.S., Ashburner, J., Smith, J., Rorden, C., 2016. The first step for neuroimaging data analysis: DICOM to NIfTI conversion. J. Neurosci. Methods 264, 47–56.

Li, Y., Liu, P., Li, Y., Fan, H., Su, P., Peng, S.-L., Park, D.C., Rodrigue, K.M., Jiang, H., Faria, A.V., Others, 2018a. ASL-MRICloud: An online tool for the processing of ASL MRI data. NMR Biomed. e4051.

Li, Y., Mao, D., Li, Z., Schär, M., Pillai, J.J., Pipe, J.G., Lu, H., 2018b. Cardiac-triggered pseudo-continuous arterial-spin-labeling: A cost-effective scheme to further enhance the reliability of arterial-spin-labeling MRI. Magn. Reson. Med. 80, 969–975.

Lorenz, K., Mildner, T., Schlumm, T., Möller, H.E., 2018. Characterization of pseudo-continuous arterial spin labeling: Simulations and experimental validation. Magn. Reson. Med. 79, 1638–1649.

Macintosh, B.J., Filippini, N., Chappell, M.A., Woolrich, M.W., Mackay, C.E., Jezzard, P., 2010. Assessment of arterial arrival times derived from multiple inversion time pulsed arterial spin labeling MRI. Magn. Reson. Med. 63, 641–647.

MacIntosh, B.J., Pattinson, K.T.S., Gallichan, D., Ahmad, I., Miller, K.L., Feinberg, D.A., Wise, R.G., Jezzard, P., 2008. Measuring the effects of remifentanil on cerebral blood flow and arterial arrival time using 3D GRASE MRI with pulsed arterial spin labelling. J. Cereb. Blood Flow Metab. 28, 1514–1522.

Madai, V.I., Martin, S.Z., von Samson-Himmelstjerna, F.C., Herzig, C.X., Mutke, M.A., Wood, C.N., Thamm, T., Zweynert, S., Bauer, M., Hetzer, S., Günther, M., Sobesky, J., 2016. Correction for Susceptibility Distortions Increases the Performance of Arterial Spin Labeling in Patients with Cerebrovascular Disease. J. Neuroimaging 26, 436–444.

Malone, I.B., Leung, K.K., Clegg, S., Barnes, J., Whitwell, J.L., Ashburner, J., Fox, N.C., Ridgway, G.R., 2015. Accurate automatic estimation of total intracranial volume: A nuisance variable with less nuisance. Neuroimage 104, 366–372.

Mato Abad, V., García-Polo, P., O’Daly, O., Hernández-Tamames, J.A., Zelaya, F., 2016. ASAP (Automatic Software for ASL Processing): A toolbox for processing Arterial Spin Labeling images. Magn. Reson. Imaging 34, 334– 344.

Maumet, C., Maurel, P., Ferré, J.-C., Bannier, E., Barillot, C., 2012. Using negative signal in mono-TI pulsed arterial spin labeling to outline pathological increases in arterial transit times. ISMRM Scientific Workshop. Perfusion MRI: Standardization, Beyond CBF & Everyday Clinical Applications 40, 42.

Maumet, C., Maurel, P., Ferre, J.C., Barillot, C., 2014. Robust estimation of the cerebral blood flow in arterial spin labelling. Magn. Reson. Imaging 32, 497–504.

Mazziotta, J., Toga, A., Evans, A., Fox, P., Lancaster, J., Zilles, K., Woods, R., Paus, T., Simpson, G., Pike, B., Holmes, C., Collins, L., Thompson, P., MacDonald, D., Iacoboni, M., Schormann, T., Amunts, K., Palomero-Gallagher, N., Geyer, S., Parsons, L., Narr, K., Kabani, N., Le Goualher, G., Boomsma, D., Cannon, T., Kawashima, R., Mazoyer, B., 2001. A probabilistic atlas and reference system for the human brain: International Consortium for Brain Mapping (ICBM). Philos. Trans. R. Soc. Lond. B Biol. Sci. 356, 1293–1322.

Melbourne, A., Toussaint, N., Owen, D., Simpson, I., Anthopoulos, T., De Vita, E., Atkinson, D., Ourselin, S., 2016. NiftyFit: a Software Package for Multi-parametric Model-Fitting of 4D Magnetic Resonance Imaging Data. Neuroinformatics 14, 319–337.

Mendrik, A.M., Vincken, K.L., Kuijf, H.J., Breeuwer, M., Bouvy, W.H., de Bresser, J., Alansary, A., de Bruijne, M., Carass, A., El-Baz, A., Jog, A., Katyal, R., Khan, A.R., van der Lijn, F., Mahmood, Q., Mukherjee, R., van Opbroek, A., Paneri, S., Pereira, S., Persson, M., Rajchl, M., Sarikaya, D., Smedby, Ö., Silva, C.A., Vrooman, H.A., Vyas, S., Wang, C., Zhao, L., Biessels, G.J., Viergever, M.A., 2015. MRBrainS Challenge: Online Evaluation Framework for Brain Image Segmentation in 3T MRI Scans. Comput. Intell. Neurosci. 2015, 813696.

MJD-ASL Webpage, https://ww2.mc.vanderbilt.edu/donahuelab/51697

Moghaddasi, L., Bezak, E., Harriss-Phillips, W., 2015. Evaluation of current clinical target volume definitions for glioblastoma using cell-based dosimetry stochastic methods. Br. J. Radiol. 88, 20150155.

Mulhollan, Z., Mutsaerts, H.-J., Petr, J., Lazar, R., Marshall, R., Asllani, I., 2018. Rethinking vascular artifacts: testing the sensitivity of ASL vascular signal as a biomarker of disease, in: ISMRM ‘18: Proceedings of the Joint Annual Meeting ISMRM - ESMRMB. p. 548.

Mutsaerts, H.J.M.M., Mirza, S.S., Petr, J., Thomas, D.L., Cash, D.M., Bocchetta, M., de Vita, E., Metcalfe, A.W.S., Shirzadi, Z., Robertson, A.D., Tartaglia, M.C., Mitchell, S.B., Black, S.E., Freedman, M., Tang-Wai, D., Keren, R., Rogaeva, E., van Swieten, J., Laforce, R., Tagliavini, F., Borroni, B., Galimberti, D., Rowe, J.B., Graff, C., Frisoni, G.B., Finger, E., Sorbi, S., de Mendonça, A., Rohrer, J.D., MacIntosh, B.J., Masellis, M., GENetic Frontotemporal dementia Initiative (GENFI), 2019. Cerebral perfusion changes in presymptomatic genetic frontotemporal dementia: a GENFI study. Brain 142, 1108–1120.

Mutsaerts, H.J.M.M., Richard, E., Heijtel, D.F.R., van Osch, M.J.P., Majoie, C.B.L.M., Nederveen, A.J., 2014a. Gray matter contamination in arterial spin labeling white matter perfusion measurements in patients with dementia. NeuroImage: Clinical 4, 139–144.

Mutsaerts, H.J.M.M., Steketee, R.M.E., Heijtel, D.F.R., Kuijer, J.P.A., Van Osch, M.J.P., Majoie, C.B.L.M., Smits, M., Nederveen, A.J., 2014b. Inter-vendor reproducibility of pseudo-continuous arterial spin labeling at 3 Tesla. PLoS One 9, e104108.

Mutsaerts, H.J.M.M., van Osch, M.J.P., Zelaya, F.O., Wang, D.J.J., Nordhøy, W., Wang, Y., Wastling, S., Fernández-Seara, M.A., Petersen, E.T., Pizzini, F.B., Fallatah, S., Hendrikse, J., Geier, O., Günther, M., Golay, X., Nederveen, A.J., Bjørnerud, A., Groote, I.R., 2015. Multi-vendor reliability of arterial spin labeling perfusion MRI using a near-identical sequence: Implications for multi-center studies. Neuroimage 113. https://doi.org/10.1016/j.neuroimage.2015.03.043

Mutsaerts, H., Petr, J., Thomas, D.L., De Vita, E., Cash, D.M., Van Osch, M.J.P., Golay, X., Groot, P.F.C.C., Ourselin, S., Van Swieten, J., Laforce, R., Jr, Tagliavini, F., Borroni, B., Galimberti, D., Rowe, J.B., Graff, C., Pizzini, F.B., Finger, E., Sorbi, S., Castelo Branco, M., Rohrer, J.D., Masellis, M., Macintosh, B.J., 2018. Comparison of arterial spin labeling registration strategies in the multi-center GENetic frontotemporal dementia initiative (GENFI). J. Magn. Reson. Imaging 47, 131–140.

Mutsaerts, H., Petr, J., Václavů, L., Van Dalen, J.W., Robertson, A.D., Caan, M.W.A., Masellis, M., Nederveen, A.J., Richard, E., Macintosh, B.J., 2017. The spatial coefficient of variation in arterial spin labeling cerebral blood flow images. J. Cereb. Blood Flow Metab. 37, 3184–3192.

Nichols, T.E., Das, S., Eickhoff, S.B., Evans, A.C., Glatard, T., Hanke, M., Kriegeskorte, N., Milham, M.P., Poldrack, R.A., Poline, J.-B., Proal, E., Thirion, B., Van Essen, D.C., White, T., Yeo, B.T.T., 2017. Best practices in data analysis and sharing in neuroimaging using MRI. Nat. Neurosci. 20, 299–303.

Oliver, R.A., 2015. Improved quantification of arterial spin labelling images using partial volume correction techniques. UCL (University College London).

Oliver-Taylor, A., Gonsalves, M., Hampshire, T., Davis, B., Daga, P., Evans, L., Bainbridge, A., Wheeler-Kingshott, C., Sokolska, M., Thornton, J., De Vita, E., Golay, X., 2017. A Calibrated Perfusion Phantom for Quality Assurance of Quantitative Arterial Spin Labelling, in: ISMRM ‘17: Proceedings of the 25th Scientific Meeting and Exhibition of International Society for Magnetic Resonance in Medicine. ISMRM, p. 681.

Pareto, D., Sastre-Garriga, J., Aymerich, F.X., Auger, C., Tintoré, M., Montalban, X., Rovira, A., 2016. Lesion filling effect in regional brain volume estimations: a study in multiple sclerosis patients with low lesion load. Neuroradiology 58, 467–474.

Petersen, E.T., Mouridsen, K., Golay, X., 2010. The QUASAR reproducibility study, Part II: Results from a multi-center Arterial Spin Labeling test-retest study. Neuroimage 49, 104–113.

Petr, J., Mutsaerts, H.J., De Vita, E., Shirzadi, Z., Cohen, S., Blokhuis, C., Pajkrt, D., Hofheinz, F., van den Hoff, J., Asllani, I., 2016. Cerebral blood flow underestimation due to volume realignments: an error induced by registration in arterial spin labeling MRI, in: European Society of Magnetic Resonance in Medicine and Biology. Vienna.

Petr, J., Mutsaerts, H.J.M.M., Vita, E.D., Steketee, R.M.E., Smits, M., Nederveen, A.J., Hofheinz, F., Van Den Hoff, J., Asllani, I., Petr, J., 2018. Effects of systematic partial volume errors on the estimation of gray matter cerebral blood flow with arterial spin labeling MRI. Magn. Reson. Mater. Phys. Biol. Med. https://doi.org/10.1007/s10334-018-0691-y

Petr, J., Platzek, I., Hofheinz, F., Mutsaerts, H.J.M.M.H., Asllani, I., Van Osch, M.J.P.M., Seidlitz, A., Krukowsky, P., Gommlich, A., Beuthien-Baumann, B., Jentsch, C., Maus, J., Troost, E.E.G.C., Baumann, M., Krause, M., van den Hoff, J., Krukowski, P., Gommlich, A., Beuthien-Baumann, B., Jentsch, C., Maus, J., Troost, E.E.G.C., Baumann, M., Krause, M., van den Hoff, J., 2018. Photon vs. proton radiochemotherapy, effects on brain tissue volume and perfusion. Radiother. Oncol. In press. https://doi.org/10.1016/j.radonc.2017.11.033

Petr, J., Schramm, G., Hofheinz, F., Langner, J., van den Hoff, J., 2013. Partial volume correction in arterial spin labeling using a Look-Locker sequence. Magn. Reson. Med. 70, 1535–1543.

Pinto, J., Chappell, M.A., Okell, T.W., Mezue, M., Segerdahl, A.R., Tracey, I., Vilela, P., Figueiredo, P., 2019. Calibration of arterial spin labeling data-potential pitfalls in post-processing. Magn. Reson. Med. https://doi.org/10.1002/mrm.28000

Pohmann, R., 2010. Accurate, localized quantification of white matter perfusion with single-voxel ASL. Magn. Reson. Med. 64, 1109–1113.

Poldrack, R.A., Baker, C.I., Durnez, J., Gorgolewski, K.J., Matthews, P.M., Munafò, M.R., Nichols, T.E., Poline, J.B., Vul, E., Yarkoni, T., 2017. Scanning the horizon: Towards transparent and reproducible neuroimaging research. Nat. Rev. Neurosci. 18, 115–126.

Power, J.D., Barnes, K.A., Snyder, A.Z., Schlaggar, B.L., Petersen, S.E., 2012. Spurious but systematic correlations in functional connectivity MRI networks arise from subject motion. Neuroimage 59, 2142–2154.

Puig, O., Vestergaard, M.B., Lindberg, U., Hansen, A.E., Ulrich, A., Andersen, F.L., Johannesen, H.H., Rostrup, E., Law, I., Larsson, H.B.W., Henriksen, O.M., 2018. Phase contrast mapping MRI measurements of global cerebral blood flow across different perfusion states – A direct comparison with O-H 2 O positron emission tomography using a hybrid PET / MR system. J. Cereb. Blood Flow Metab. 0, 1–11.

Regulation, G.D.P., 2016. Regulation (EU) 2016/679 of the European Parliament and of the Council of 27 April 2016 on the protection of natural persons with regard to the processing of personal data and on the free movement of such data, and repealing Directive 95/46. Official Journal of the European Union (OJ) 59, 294.

Restom, K., Behzadi, Y., Liu, T.T., 2006. Physiological noise reduction for arterial spin labeling functional MRI. Neuroimage 31, 1104–1115.

Ripollés, P., Marco-Pallarés, J., de Diego-Balaguer, R., Miró, J., Falip, M., Juncadella, M., Rubio, F., Rodriguez-Fornells, A., 2012. Analysis of automated methods for spatial normalization of lesioned brains. Neuroimage 60, 1296–1306.

Ritchie, C.W., Molinuevo, J.L., Truyen, L., Satlin, A., Van der Geyten, S., Lovestone, S., 2016. Development of interventions for the secondary prevention of Alzheimer’s dementia: the European Prevention of Alzheimer’s Dementia (EPAD) project. The Lancet Psychiatry 3, 179–186.

Robertson, A.D., Matta, G., Basile, V.S., Black, S.E., Macgowan, C.K., Detre, J.A., MacIntosh, B.J., 2017. Temporal and Spatial Variances in Arterial Spin-Labeling Are Inversely Related to Large-Artery Blood Velocity. AJNR Am. J. Neuroradiol. 38, 1555–1561.

Sayer, N., 2014. Google Code Archive-Long-term storage for Google Code Project Hosting. XP055260798, Retrieved from the Internet [retrieved on 20160323].

Schmid, S., Heijtel, D.F.R., Mutsaerts, H.J.M.M., Boellaard, R., Lammertsma, A.A., Nederveen, A.J., Van Osch, M.J.P., Van Osch, M.J.P., 2015. Comparison of velocity- and acceleration-selective arterial spin labeling with [15O]H2O positron emission tomography. J. Cereb. Blood Flow Metab. 35, 1–8.

Schmidt, P., Gaser, C., Arsic, M., Buck, D., Forschler, A., Berthele, A., Hoshi, M., Ilg, R., Schmid, V.J., Zimmer, C., Hemmer, B., Muhlau, M., 2012. An automated tool for detection of FLAIR-hyperintense white-matter lesions in Multiple Sclerosis. Neuroimage 59, 3774–3783.

Shin, D.D., Ozyurt, I.B., Brown, G.G., Fennema-notestine, C., Liu, T.T., 2016. NeuroImage The Cerebral Blood Flow Biomedical Informatics Research Network (CBFBIRN) data repository. Neuroimage 124, 1202–1207.

Shirzadi, Z., Crane, D.E., Robertson, A.D., Maralani, P.J., Aviv, R.I., Chappell, M.A., Goldstein, B.I., Black, S.E., MacIntosh, B.J., 2015. Automated removal of spurious intermediate cerebral blood flow volumes improves image quality among older patients: A clinical arterial spin labeling investigation. J. Magn. Reson. Imaging 42, 1377–1385.

Shirzadi, Z., Stefanovic, B., Chappell, M.A., Ramirez, J., Schwindt, G., Masellis, M., Black, S.E., MacIntosh, B.J., 2018. Enhancement of automated blood flow estimates (ENABLE) from arterial spin-labeled MRI. J. Magn. Reson. Imaging 47, 647–655.

Smith, S.M., 2002. Fast robust automated brain extraction. Hum. Brain Mapp. 17, 143–155.

Spann, S.M., Kazimierski, K.S., Aigner, C.S., Kraiger, M., Bredies, K., Stollberger, R., 2017. Spatio-temporal TGV denoising for ASL perfusion imaging. Neuroimage 157, 81–96.

Steketee, R.M.E., Bron, E.E., Meijboom, R., Houston, G.C., Klein, S., Mutsaerts, H.J.M.M., Mendez Orellana, C.P., de Jong, F.J., van Swieten, J.C., van der Lugt, A., Smits, M., 2016. Early-stage differentiation between presenile Alzheimer’s disease and frontotemporal dementia using arterial spin labeling MRI. Eur. Radiol. 26, 244–253.

Tan, H., Maldjian, J.A., Pollock, J.M., Burdette, J.H., Yang, L.Y., Deibler, A.R., Kraft, R.A., 2009. A fast, effective filtering method for improving clinical pulsed arterial spin labeling MRI. J. Magn. Reson. Imaging 29, 1134– 1139.

Ten Kate, M., Ingala, S., Schwarz, A.J., Fox, N.C., Chételat, G., van Berckel, B.N.M., Ewers, M., Foley, C., Gispert, J.D., Hill, D., Irizarry, M.C., Lammertsma, A.A., Molinuevo, J.L., Ritchie, C., Scheltens, P., Schmidt, M.E., Visser, P.J., Waldman, A., Wardlaw, J., Haller, S., Barkhof, F., 2018. Secondary prevention of Alzheimer’s dementia: neuroimaging contributions. Alzheimers. Res. Ther. 10, 112.

Tohka, J., Zijdenbos, A., Evans, A., 2004. Fast and robust parameter estimation for statistical partial volume models in brain MRI. Neuroimage 23, 84–97.

Václavů, L., Meynart, B.N., Mutsaerts, H.J., Petersen, E.T., Majoie, C.B., VanBavel, E.T., Wood, J.C., Nederveen, A.J., Biemond, B.J., 2018. Hemodynamic provocation with acetazolamide shows impaired cerebrovascular reserve in adults with sickle cell disease. Haematologica. https://doi.org/10.3324/haematol.2018.206094

Vaclavu, L., van der Land, V., Heijtel, D.F.R., van Osch, M.J.P., Cnossen, M.H., Majoie, C.B.L.M., Bush, A., Wood, J.C.C., Fijnvandraat, K.J.J., Mutsaerts, H.J.M.M., Nederveen, A.J., 2016. In vivo T1 of blood measurements in children with sickle cell disease improve cerebral blood flow quantification from arterial spin-labeling MRI. AJNR Am. J. Neuroradiol. 37, 1727–1732.

Vidorreta, M., Balteau, E., Wang, Z., De Vita, E., Pastor, M. a., Thomas, D.L., Detre, J. a., Fernández-Seara, M. a., 2014. Evaluation of segmented 3D acquisition schemes for whole-brain high-resolution arterial spin labeling at 3 T. NMR Biomed. 27, 1387–1396.

Vidorreta, M., Wang, Z., Rodríguez, I., Pastor, M.A., Detre, J.A., Fernández-Seara, M.A., 2013. Comparison of 2D and 3D single-shot ASL perfusion fMRI sequences. Neuroimage 66, 662–671.

Wang, D.J.J., Chen, Y., Fernández-Seara, M.A., Detre, J.A., Fernandez-Seara, M.A., Detre, J.A., Fernández-Seara, M.A., Detre, J.A., 2011. Potentials and challenges for arterial spin labeling in pharmacological magnetic resonance imaging. J. Pharmacol. Exp. Ther. 337, 359–366.

Wang, Z., 2014. Support vector machine learning-based cerebral blood flow quantification for arterial spin labeling MRI. Hum. Brain Mapp. 35, 2869–2875.

Wang, Z., 2012. Improving cerebral blood flow quantification for arterial spin labeled perfusion MRI by removing residual motion artifacts and global signal fluctuations. Magn. Reson. Imaging 30, 1409–1415.

Wang, Z., Aguirre, G.K., Rao, H., Wang, J., Fernández-Seara, M.A., Childress, A.R., Detre, J.A., 2008. Empirical optimization of ASL data analysis using an ASL data processing toolbox: ASLtbx. Magn Reson.Imaging 26, 261–269.

Warmuth, C., Günther, M., Zimmer, C., 2003. Quantification of blood flow in brain tumors: comparison of arterial spin labeling and dynamic susceptibility-weighted contrast-enhanced MR imaging. Radiology 228, 523–532.

Wells, J. a., Thomas, D.L., King, M.D., Connelly, A., Lythgoe, M.F., Calamante, F., 2010. Reduction of errors in ASL cerebral perfusion and arterial transit time maps using image de-noising. MRM 64, 715–724.

Wilke, M., Altaye, M., Holland, S.K., CMIND Authorship Consortium, 2017. CerebroMatic: A Versatile Toolbox for Spline-Based MRI Template Creation. Front. Comput. Neurosci. 11, 5.

Ye, F.Q., Frank, J.A., Weinberger, D.R., McLaughlin, A.C., 2000. Noise reduction in 3D perfusion imaging by attenuating the static signal in arterial spin tagging (ASSIST). Magn. Reson. Med. 44, 92–100.

Zhang, K., Herzog, H., Mauler, J., Filss, C., Okell, T.W., Kops, E.R., Tellmann, L., Fischer, T., Brocke, B., Sturm, W., Coenen, H.H., Shah, N.J., 2014. Comparison of cerebral blood flow acquired by simultaneous [15O]water positron emission tomography and arterial spin labeling magnetic resonance imaging. JCBFM 34, 1373–1380.

Zhao, M.Y., Mezue, M., Segerdahl, A.R., Okell, T.W., Tracey, I., Xiao, Y., Chappell, M.A., 2017. A systematic study of the sensitivity of partial volume correction methods for the quantification of perfusion from pseudo-continuous arterial spin labeling MRI. Neuroimage 162, 384–397.

Zhu, H., Zhang, J., Wang, Z., 2018. Arterial spin labeling perfusion MRI signal denoising using robust principal component analysis. J. Neurosci. Methods 295, 10–19.

