## Supplemental Tables and Figures for "ExploreASL: an image processing pipeline for multi-center ASL perfusion MRI studies"

### Supplementary Material

| Example sequence | 1 | 2 | 3 | 4 | 5 | 6 |
|---|---|---|---|---|---|---|
| Study name | IN-OT | VESPA | Sleep | PhilipsResearch | AAS | GIFMI Mood |
| Session included | Baseline | Test and retest | All | Test and retest | Controls | Pilot (n=13) |
| Scanning location | Institute of Psychiatry, King's College London, United Kingdom | Erasmus Rotterdam Medical Center, Netherlands | Oslo University Hospital, Norway | Best, Netherlands | Oslo University Hospital, Norway | Ghent University, Belgium |
| Sample size = subjects * measurements | 32 | 22*2=44 | 39*3=117 | 34*2=68 | 68 | 13*19=247 |
| Age (years) | 25.0 ± 3.1 | 22.6 ± 2.1 | 22.1 ± 2.5 | 57.8 ± 17.0 | 31.4 ± 9.1 | 23.2 ± 2.8 |
| Sex (n Male) | 32 (100%) | 9 (40.9%) | 39 (100%) | 20 (58.8%) | 39 (100%) | 4 (38.5%) |
| Scanner | GE Signa HDx 3T | GE MR750 3T | Philips Achieva 3T | Philips Achieva 3T | Siemens Skyra 3T | Siemens Trio 3T |
| Sequence | 3D spiral PCASL (WIP) | 3D spiral PCASL | 2D EPI PCASL | 3D GRASE PCASL | 2D EPI PCASL | 3D GRASE PCASL |
| TE/TR (ms) | 4/5500 | 10.5/4600 | 11/4400 | 12/4100 | 12/4360 | 17.2/4000 |
| PLD range (ms) | 1525 | 1525 | 1800-2640 | 1800 | 1771-2759 | 2000 |
| Labeling duration (ms) | 1450 | 1450 | 1800 | 1800 | 1800 | 1500 |
| Voxel size (mm X Y Z) | 3.8 x 3.8 x 4.0 | 3.8 x 3.8 x 4.0 | 3.5 x 3.5 x 6.0 | 3.75 x 3.75 x 6.0 | 3.5 x 3.5 x 6.0 | 4.0 x 4.0 x 5.0 |
| Background suppression pulses (n) | 5 | 5 | 2 | 4 | 0 | 5 |

Supplementary Table 1 for Figure 2. Studies were 1) IN-OT = INtranasal OxyTocin trial (Paloyelis et al., 2016), 2) VESPA =VEndor SPecific features and reproducibility of Asl (Mutsaerts et al., 2014b) , 3) Sleep (Elvsåshagen et al., 2018), 4) PhilipsResearch (Baas et al., 2018), 5) AAS = Anabolic Androgenic-Steroid Research Study (Bjørnebekk et al., 2017), 6) GIFMI Mood = Ghent Institute for Functional and Metabolic Imaging, Mood study (Clement P et al., 2016). PLD ranges include the PLD increase with ascending slices for 2D acquisitions. EPI = echo-planar imaging, GE = General Electric, GRASE = GRadient echo And Spin Echo, PCASL = pseudo-continuous ASL, PLD = post-labeling delay, TE = echo time, TR = repetition time, WIP = work-in-progress sequence.

| Parameter name | Unit | Brief explanation |
|---|---|---|
| **Structural** | | |
| CSF_vol_mL | mL | CSF volume as segmented by CAT12 (Gaser et al., 2009) |
| GM_ICV_Ratio | | Ratio of GM volume to ICV volume |
| GM_vol_mL | mL | GM volume as segmented by CAT12 (Gaser et al., 2009) |
| ICV_vol_mL | mL | GM volume + WM volume + CSF volume |
| IQR_Perc | % | Image quality rate (Dahnke et al., 2015) as computed by CAT12 (Gaser et al., 2009): combination of noise, inhomogeneity, and resolution ratings. Currently tested for 3D T1w only (higher = better) |
| LR_flip_YesNo | true/false | Indicates left-right orientation flipping through registration (by comparing determinants between current and original orientation matrices) |
| WM_vol_mL | mL | WM volume as segmented by CAT12 (Gaser et al., 2009) |
| WMH_n | n | Number of non-connected segmented WMH |
| WMH_vol_mL | mL | Total lesion volume of WMH segmentation (allowing either probability map or binary mask) |
| **Structural QC parameters** | | **Repeated for T1w and FLAIR, parameters are taken from the Preprocessed Connectome Project Quality ASsurance Protocol (QAP)** ("PCP Quality Assessment Protocol," n.d.)**, adapted from the SPM Univariate Plus toolbox** (Pernet et al., 2019)**.** A small WM region is used as noise reference instead of a background region outside the brain, as background noise may not be reliable in case of parallel imaging (Magnotta et al., 2006). GM and WM masks are made mutually exclusive by thresholding. |
| CNR_GM_WM_Ratio | | abs(Mean GM - Mean WM)/(SD WMref)$^2$ (higher = better) (Magnotta et al., 2006) |
| EFC_bits | bits | Shannon entropy of voxel intensities proportional to maximum possible entropy for similarly sized image, indicating ghosting and head motion-induced blurring (lower = better) (Atkinson et al., 1997) |
| FBER_WMref_Ratio | | (SD WholeBrain)$^2$ / (SD WMref)$^2$ (higher = better) |
| Mean_AI_Perc | % | Mean of voxel-wise asymmetry index (AI), where AI = (L-R)/(0.5*[L+R]) (Kurth et al., 2015) (lower = better) |
| SD_AI_Perc | % | Distribution of voxel-wise asymmetry index (AI), where AI = (L-R)/(0.5*[L+R]) (Kurth et al., 2015) (lower = better) |
| SD_WMref | mL/100g/min | SD in the WMref region |
| SNR_GM_Ratio | | Mean GM / SD_WMref(higher = better) (Magnotta et al., 2006) |
| WMref_vol_mL | mL | Volume of noise reference region within the WM (WMref) |
| WMref_vol_Perc | % | Percentage of WMref volume to the total WM volume |
| **ASL** | | |
| CBF_GM_Median_mL100gmin | mL/100g/min | Median CBF within the individual pGM>0.5 ROI |
| CBF_GM_WM_Ratio | | GM-WM CBF Ratio, calculated as CBF_GM_median_mL100gmin/ CBF_WM_median_mL100gmin |
| CBF_WM_Median_mL100gmin | mL/100g/min | Median CBF within the individual pWM>0.5 ROI |
| Matrix_[X Y Z N] | n | Number of data elements (voxels) in each dimension |
| MotionExcl_Perc | % | Percentage of control-label pairs excluded by ENABLE (Shirzadi et al., 2015) |
| MotionMean_mm | mm | Mean motion (temporal difference NDV) as computed by SPM12 realign, acknowledging control-label zigzag (Wang et al., 2008). NDV is computed |

| | | |
|---|---|---|
| | | as RMS of translations and rotations, assuming a brain radius of 50 mm (Power et al., 2012) |
| RescaleSlope | a.u. | Scale factor used by Philips for window-leveling on scanner console |
| RigidBody2Anat_mm | mm | Registration offset between ASL and 3D-T1w anatomical scans (NDV) |
| ScaleSlope | a.u. | Scale factor used by Philips for storing floating point information in non-floating point data format |
| SpatialCoV_GM_Perc | % | Spatial CoV (*100%) within the individual pGM>0.5 ROI |
| SpatialCoV_WM_Perc | % | Spatial CoV (*100%) within the individual pWM>0.5 ROI |
| TE_ms | ms | TE from DICOM header |
| TR_ms | ms | TR from DICOM header |
| VoxelSize_[X Y Z]_mm | mm | Size of a voxel |
| **ASL QC parameters** | | **QC parameters adapted from the fmRI QC parameters** (Pernet, 2019)**.** pGM, pWM and pCSF are thresholded above 50%. As noise reference region, an eroded WM region (WMref) (Mutsaerts et al., 2014a) is used instead of a background region, as background noise may not be reliable in case of parallel imaging (Magnotta et al., 2006). |
| tSNR_CSF_Ratio | | Mean / temporal SD within total CSF (Liu, 2017) |
| tSNR_GM_Ratio | | Mean / temporal SD within total GM (Liu, 2017) |
| tSNR_GMWM_Ratio | | Mean / temporal SD within the parenchyma (GM+WM) |
| tSNR_GMWM_WMref_Ratio | | Mean within parenchyma / tSD within WMref |
| tSNR_Physio2Thermal_Ratio | | tSNR_GMWM / tSNR_GMWM_WMref (Wald et al., 2017) |
| tSNR_Slope_Corr | | Slope of temporal SNR as function of GM partial volume |
| tSNR_WM_Ratio | | Mean / temporal SD within total WM (Liu et al., 2017) |
| tSNR_WMref_Ratio | | Mean / temporal SD within WMref |
| **QC_diff_template** | | **Experimental QC parameters that compare an image with a template** computed after 8 mm FWHM Gaussian smoothing of individual image within a whole-brain mask |
| AI_Perc | % | Mean voxel-wise asymmetry index, computed as (L-R)/(0.5*[L+R]) (Kurth et al., 2015) (lower = better) |
| Mean_SSIM_Perc | % | Mean voxel-wise structural similarity index (Wang et al., 2004) (higher = better) |
| nRMSE_Perc | % | nRMSE with ASL template (lower = better) |
| PeakSNR_Ratio | | Assuming an individual ASL image can be interpreted as a noisy version of an ASL template, this parameter is the dynamic range compared to the mean squared difference of the individual and template ASL images. Dynamic range is calculated here as MaxIntensity-MinIntensity (higher = better) |
| RMSE_Perc | % | RMSE with ASL template (lower = better) |

Supplementary Table 2. Legend of ExploreASL automatic quality control (QC), of which a visual example is shown in Figure 3. CAT12 = Computational Anatomy Toolbox, CBF = cerebral blood flow, CoV = coefficient of variation, CNR = contrast-to-noise ratio, CSF = cerebrospinal fluid, EFC = entropy focus criterion, FBER = foreground to background energy ratio, FLAIR = FLuid Attenuation Inversion Recovery, GM = gray matter, ICV = intracranial volume, L = left, NDV = net displacement vector, QC = quality control, Perc = percentage, R = right, ROI = region-of-interest, RMSE = root-mean-square error, PSNR = peak SNR, SSIM = structural similarity, SNR = signal-to-noise ratio, SPM = statistical parametric mapping, SPM U+ = SPM Univariate Plus, TE = echo time, TR = repetition time, WM = white matter, WMH = WM hyperintensity, TLV = total lesion volume, tSD = temporal SD, tSNR = temporal SNR.

| **Example study** | **1** | **2** | **3** |
|---|---|---|---|
| Study name | Neurological, cOgnitive and VIsual performance in perinatally HIV-infected ChildrEn (NOVICE) (Blokhuis et al., 2017) | Healthy adults (Circadian/Sleep deprivation study) (Elvsåshagen et al., 2018) | European Prevention of Alzheimer's Dementia (EPAD) (Ritchie et al., 2016; Ten Kate et al., 2018) |
| Session included | Baseline, single-center | Baseline, single-center | Baseline, single-center |
| Group included | perinatally HIV-infected children | Healthy adult males | Older adults with mild cognitive complaints |
| Scanning location | Amsterdam UMC | University of Oslo | Amsterdam UMC |
| Sample size (n) | 28 | 38 | 84 |
| Age (years) | 13.5 ± 2.1 | 22.1 ± 2.5 | 67.7 ± 7.1 |
| Sex (n Male (%)) | 15 (54%) | 38 (100%) | 38 (45.2%) |
| Scanner | Philips 3T Ingenia | Philips 3T Achieva | Philips 3T Achieva |
| Sequence | 2D EPI PCASL | 2D EPI PCASL | 2D EPI PCASL |
| TE/TR (ms) | 14/4000 | 11/4400 | 10.4/4800 |
| PLD range (ms) | 1525-2230 | 1800-2640 | 2025-3310 |
| Labeling duration (ms) | 1650 | 1800 | 1650 |
| Acquisition voxel size ($mm^3$) | [3.0 3.0 6.6] | [3.5 3.5 6.0] | [3.5 3.5 4.5] |

Supplementary Table 3. EPI = echo-planar imaging, HIV = Human Immunodeficiency Virus, PCASL = pseudo-continuous ASL, PLD = post-labeling delay, TE = echo time, TR = repetition time, UMC = University Medical Center

| | NOVICE | | Sleep | | EPAD | |
|---|---|---|---|---|---|---|
| | pre | post | pre | post | pre | post |
| 2.1 WMH (%) | 0.58 | 1.27 | 0.00 | 0.00 | 0.83 | 0.45 |
| 2.2 pGM (%) | 0.22 | 1.12 | 0.00 | 0.00 | 0.31 | 0.25 |
| 2.2 T1w position (mm) | 0.05 | 0.05 | 0.00 | 0.00 | 0.31 | 0.02 |
| 3.2 ASL position (mm) | 0.19 | 0.34 | 0.42 | 0.15 | 0.75 | 0.52 |
| 3.4 pGM (%) | 0.85 | 1.79 | 0.73 | 0.47 | 2.05 | 1.42 |
| 3.5 CBF (%) | 0.06 | 0.30 | 0.27 | 0.22 | 0.17 | 0.09 |
| 3.7 GM CBF (%) | 0.52 | 0.95 | 1.22 | 0.57 | 2.72 | 1.77 |

Supplementary Table 4. Showing reproducibility across two centers with different OSes and MATLAB versions (Linux-2018b vs Windows-2015a). Reproducibility rows follow the pipeline sections as described in the Theory section (in brackets). ASL = arterial spin labeling, CBF = cerebral blood flow, GM CBF = CBF in GM after PV correction, GM = gray matter, PV = partial volume, WMH = white matter hyperintensities.

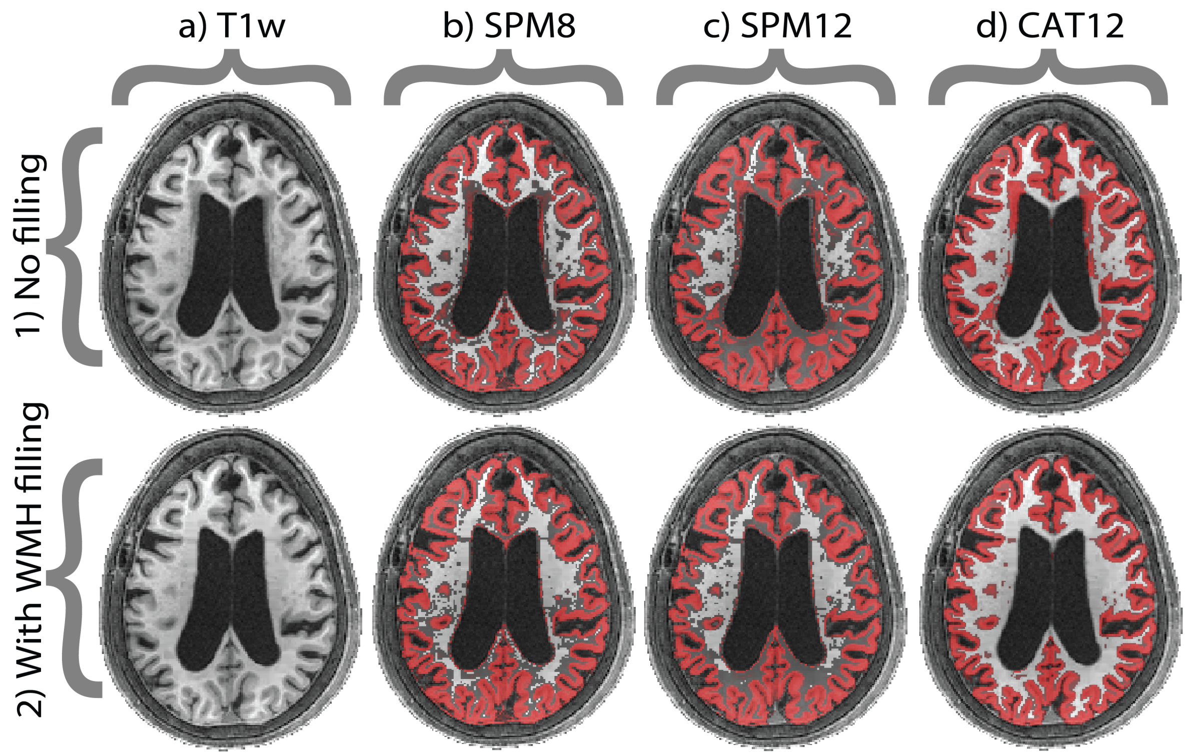


Supplementary Figure 1. Comparison between SPM-based segmentation algorithms (b-d), without (1) or with (2) being preceded by WMH lesion filling. These differences are more pronounced in subjects with large atrophy and/or lesions (such as the example shown in this Figure). Gray matter segmentation is projected in red over the T1w image. By showing the full range of the probabilities, there is a clear difference between (d) CAT12 - which segments partial volume - and the SPM methods (b-c). CAT = Computational Anatomical Toolbox, SPM = Statistical Parametric Mapping, T1w = T1-weighted, WMH = White Matter Hyperintensity. Example data are from the Singapore Memory Clinic Study (Ferro et al., 2019).

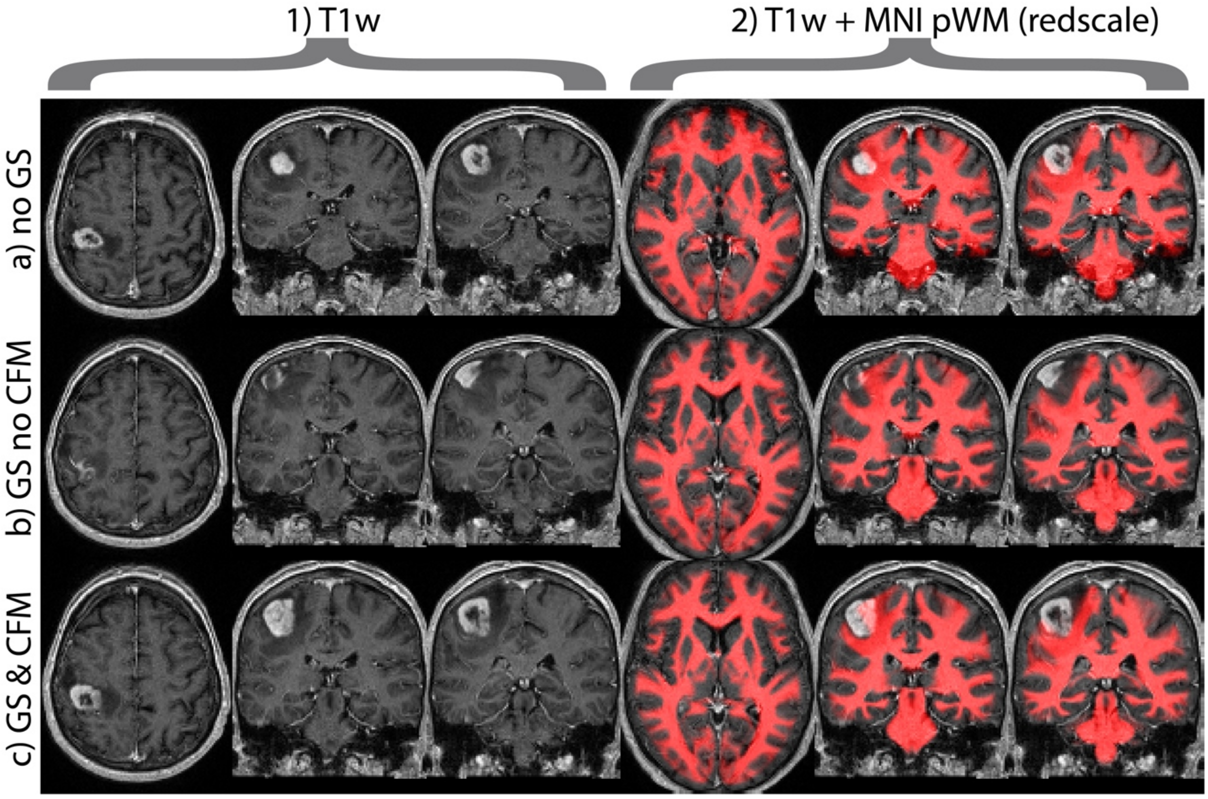


Supplementary Figure 2. Columns show a transversal and two coronal T1w slices (1) and T1w slices with MNI template WM segmentation overlaid in red (2). With linear registration only, the tumor is not distorted (a1) but the registration to MNI is not sufficient for the normal tissue (a2). Non-linear registration improves the alignment for the normal tissue (b2) - compare the position of the WM template around the ventricles and in the temporal lobes - but it pushes the tumor outside of the brain (b1). Non-linear registration with CFM allows the same desired non-linear registration effect for the normal tissue (c2) while restricting unrealistic deformations of the tumor (c1). CFM = cost-function masking. GM = gray matter, GS = geodesic shooting (non-linear registration), MNI = Montreal Neurological Institute standard space, WM = white matter. Example data from the PICTURE study (Visser et al., 2019).

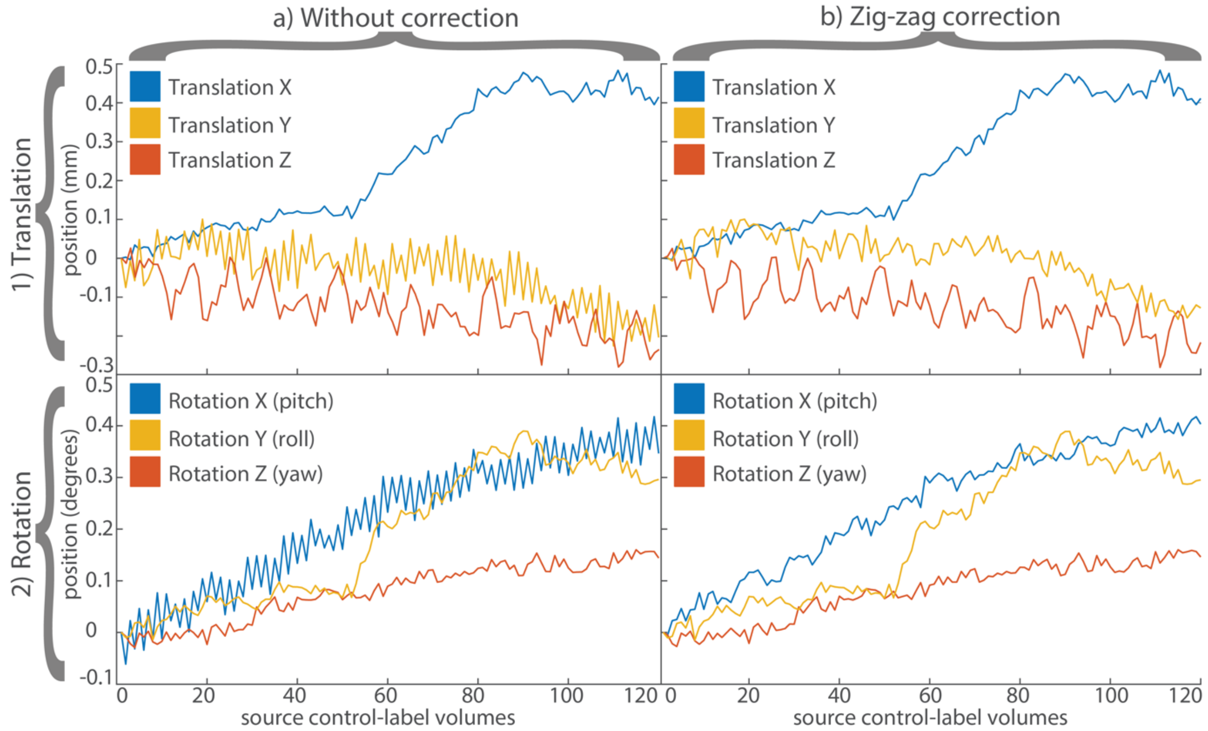


Supplementary Figure 3. Translations in mm (1) and rotations in degrees (2) without (a) and with (b) zig-zag regression, for motion estimation of a single healthy control individual with 2D EPI PCASL 60 control-label pairs. The zig-zag regression effect is visible in the Z-translation (1a, yellow) and pitch rotation (2a, blue), which both describe through-slice motion without changing the left-right symmetry. This effect is visibly removed in the respective curves after zig-zag regression (b). Data are taken from the Sleep study (Elvsåshagen et al., 2018).

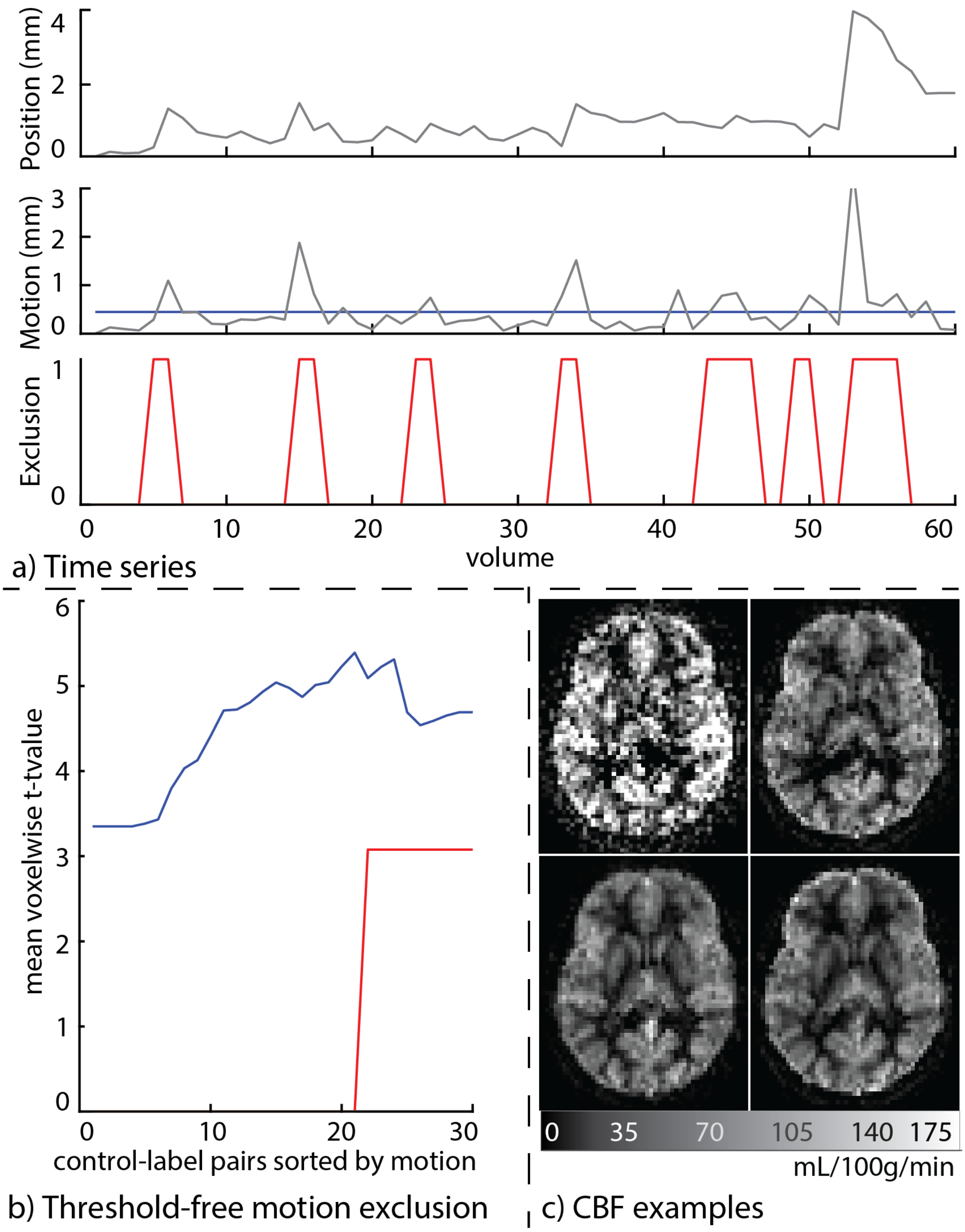


Supplementary Figure 4. The ability of ENABLE to detect excessive motion and artifacts for exclusion on a pediatric dataset: a) Position, which is the net displacement vector: the RMS of the three translations and rotations relative to the first volume. Motion is the difference in the position in two succeeding volumes. The blue line depicts the mean motion. b) The cumulative increase in t-values with the addition of more control-label pairs, sorted by ascending motion, is shown as a blue curve. The t-values decrease when volumes with too high motion are added. These volumes — 30% of the total number of volumes — are excluded by ENABLE (red line in a) and b)). c) From left to right, upper to lower: a single control-label subtraction, 5 pairs averaged, the optimum pairs averaged according to ENABLE, all pairs averaged. Note the hyperintensity rim in the 4th image, that is not apparent in the 3rd image. NDV = net displacement vector; RMS = root mean square. RMS = root mean square. ENABLE = ENhancement of Automated BLood flow Estimates (Shirzadi et al., 2015). Data are taken from the NOVICE study (Blokhuis et al., 2017).

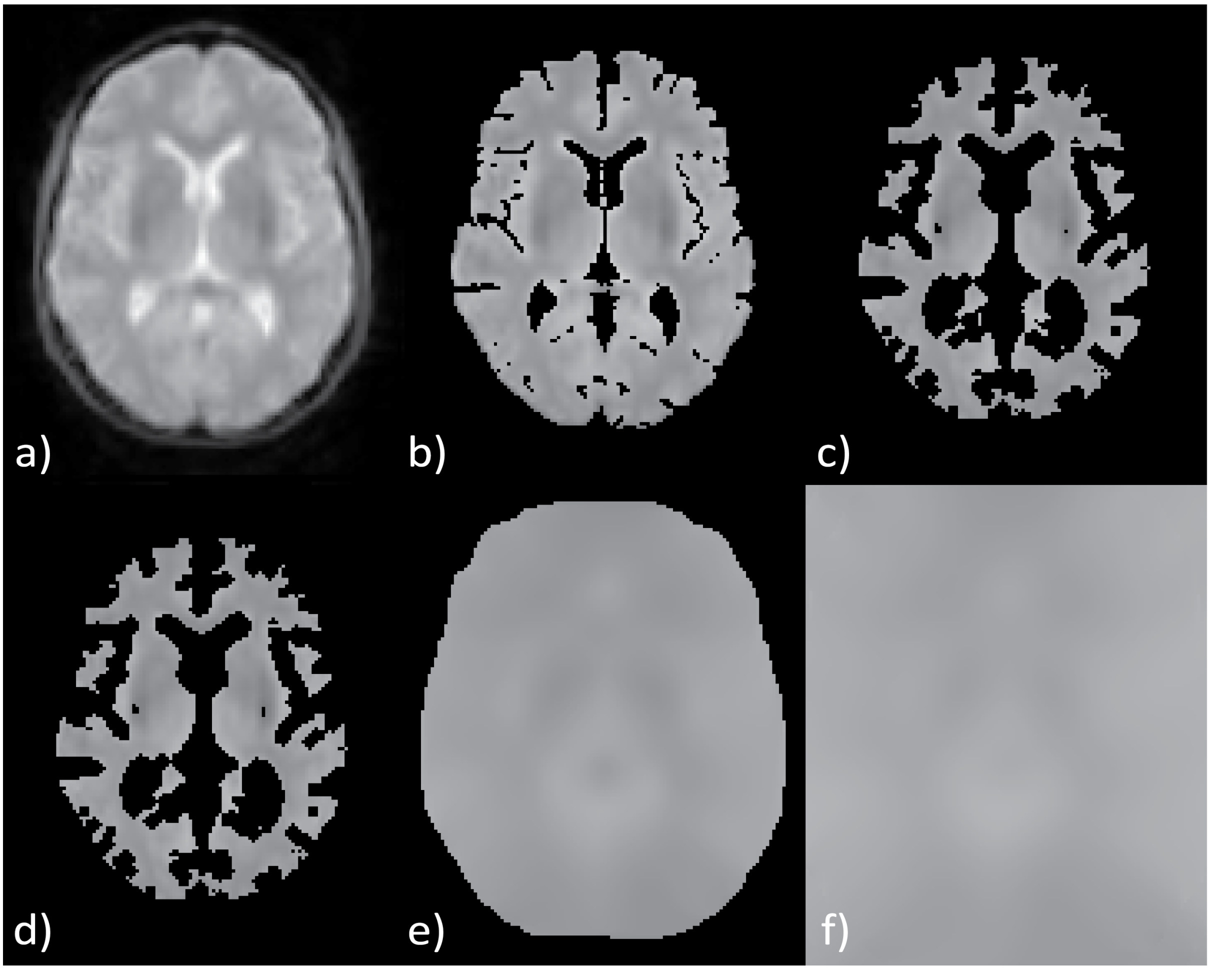


Supplementary Figure 5. ExploreASL M0 image processing steps shown for one transverse slice. The original M0 image (a) is masked with a (pGM+pWM)>50% mask (b), eroded with a two-voxel sphere to limit the influence of the ventricular and extracranial signal (c), and thresholded to exclude significantly high (i.e. median + 3*mean absolute deviation (MAD)) border region values (d). This masked M0 image is smoothed with a 16x16x16 mm full-width-half-maximum Gaussian filter (Mutsaerts et al., 2018) (e), after which the signal at the border is smoothly extrapolated until the full image is filled (f). Whereas the masking avoids mixing with cerebrospinal fluid or extracranial signal, the extrapolation avoids M0 division artifacts (see Supplementary Figure 6 created using data from the same subject). Data from the EPAD study (Ritchie et al., 2016; Ten Kate et al., 2018).

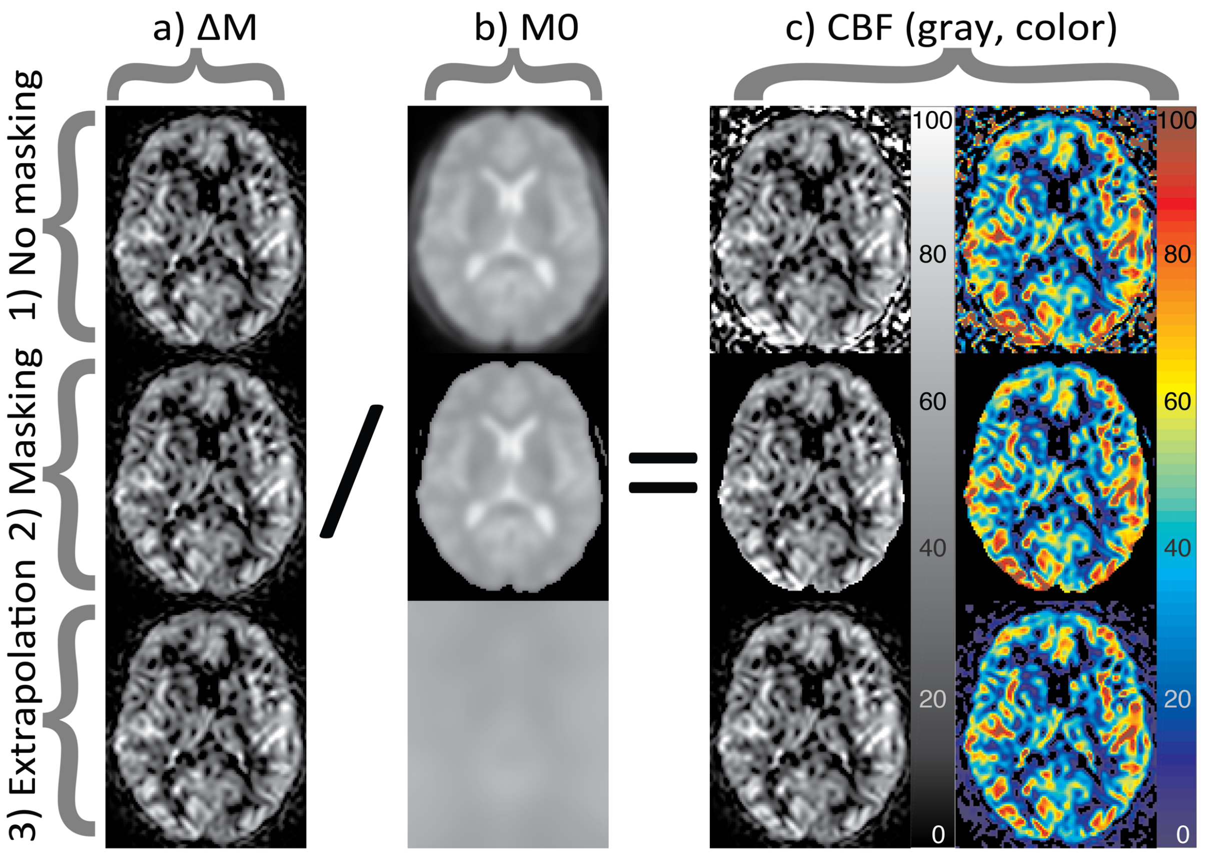


Supplementary Figure 6. M0 division artifacts in CBF without (1) and with M0 masking (2). ExploreASL aims to reduce the division artifacts by using M0 extrapolation instead of masking (3). Identical ΔMs (a) are divided by different M0s (b) resulting in different CBF images (c, in ml/min/100g). (a) and (b) show the typically recommended/traditional approach (Alsop et al., 2015): i.e. mild smoothing and masking. Note that the division artifacts are reduced but not completely eliminated by masking. Moreover, masking can remove parts of the CBF image which becomes problematic for smooth 3D ASL acquisitions (data not shown). Also note that the masking prohibits to analyze extracranial artifacts, which could be useful for quality control. The color-scaled CBF image (c) shows the subtle differences between the ExploreASL method and previous methods: in the ExploreASL method, the cerebrospinal fluid and extracranial values were masked out before smoothing. Note the differences in asymmetry in the subcortical perfusion, as well as the differences in posterior perfusion. CBF = cerebral blood flow, ΔM= perfusion weighted difference image. Figure was created using the data from the same EPAD participant as in Supplementary Figure 5 (Ritchie et al., 2016; Ten Kate et al., 2018).

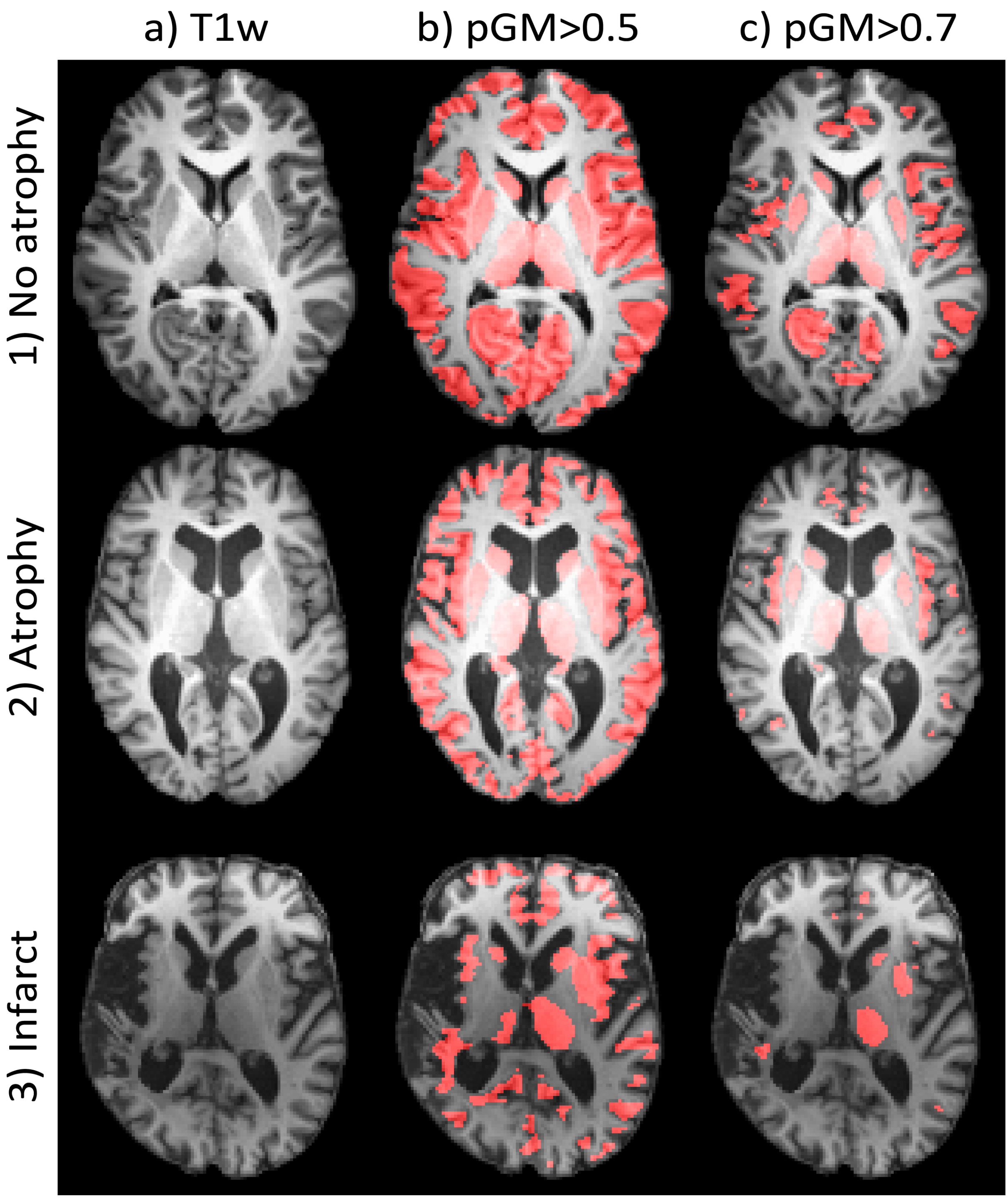


Supplementary Figure 7. Showing the necessity of partial volume correction in ASL for three subjects: 1) a healthy adult (Sleep dataset (Elvsåshagen et al., 2018)), 2) an older adult with atrophy (EPAD dataset (Ritchie et al., 2016; Ten Kate et al., 2018)), and 3) an older adult with a unilateral infarct (Singapore Memory Clinic Study (Ferro et al., 2019)). a) Shows the native space T1-weighted (T1w) images, b, c) show the T1w images overlaid, in red, with the GM segmentations smoothed to the ASL resolution (3x3x7 $mm^3$ here) and thresholded at b) 50% and c) 70%.The pGM threshold of 70% is typically used for calculating the mean GM CBF with and without PVC. These images show that, especially in clinical cases and thin GM regions, only a fraction of ASL voxels contain sufficient GM to pass the thresholding, thus introducing a spatial bias in the mean CBF calculation. Use of PVC with a 50% threshold is thus recommended to have reasonable spatial coverage while avoiding PV effects. Note that these effects can be even stronger for ASL readouts with lower effective resolution (e.g. 3D readouts). GM = gray matter, pGM = partial volume GM. PVC = partial volume correction.

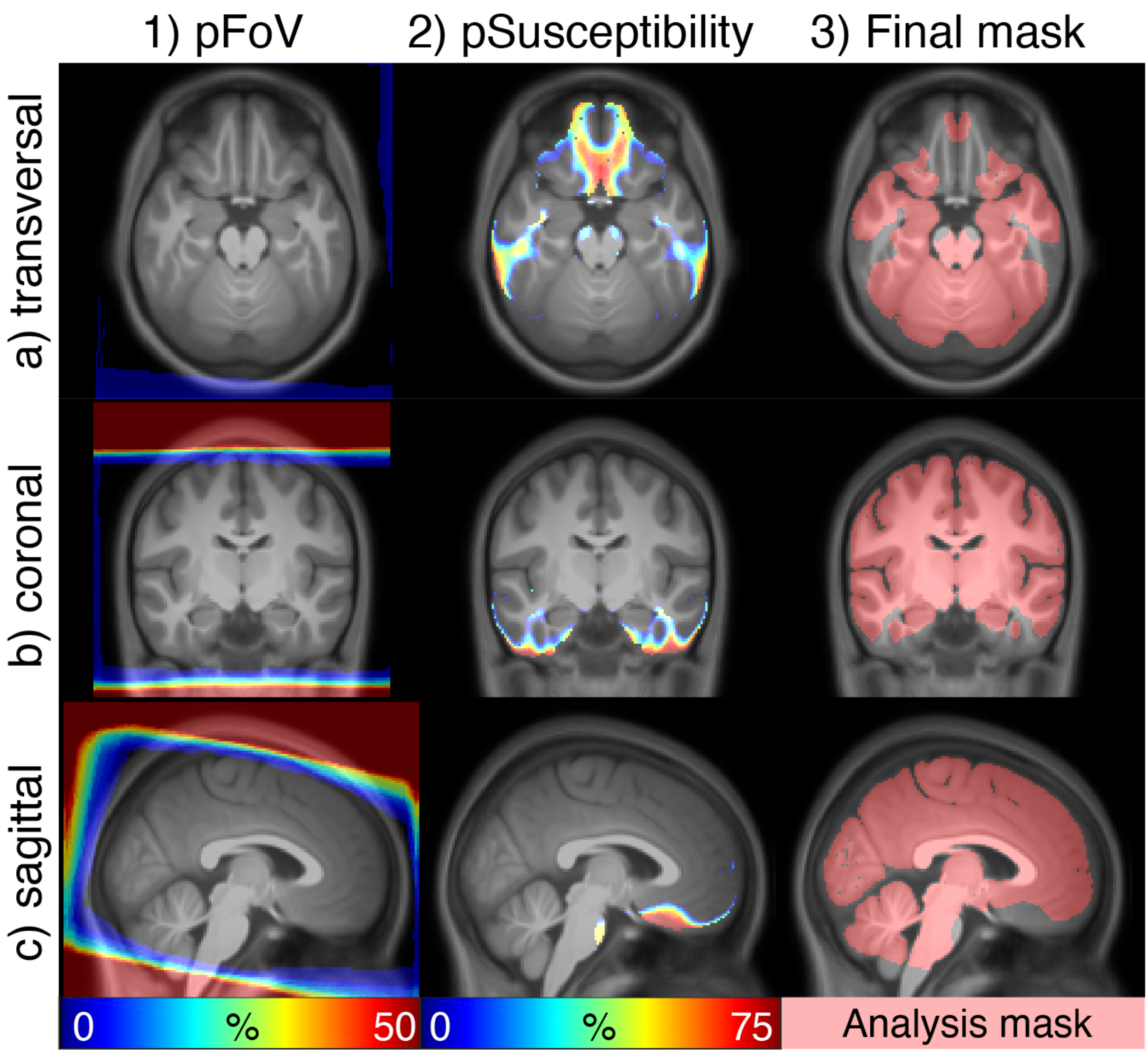


Supplementary Figure 8. Overview of the creation of the population-based analysis mask in standard space overlaid over the population-average T1w image. 1) and 2) Show the voxel-wise percentage of being outside the FoV mask (1) and susceptibility mask (2). Note that the subjects' heads were positioned similarly in the same head coil resulting in minor FoV differences (1). Susceptibility artifact masks vary within the predefined regions surrounding the skull air cavities (2). 3) The resulting group analysis mask in red. p = probability, FoV = field-of-view. Data from the NOVICE dataset (Blokhuis et al., 2017)

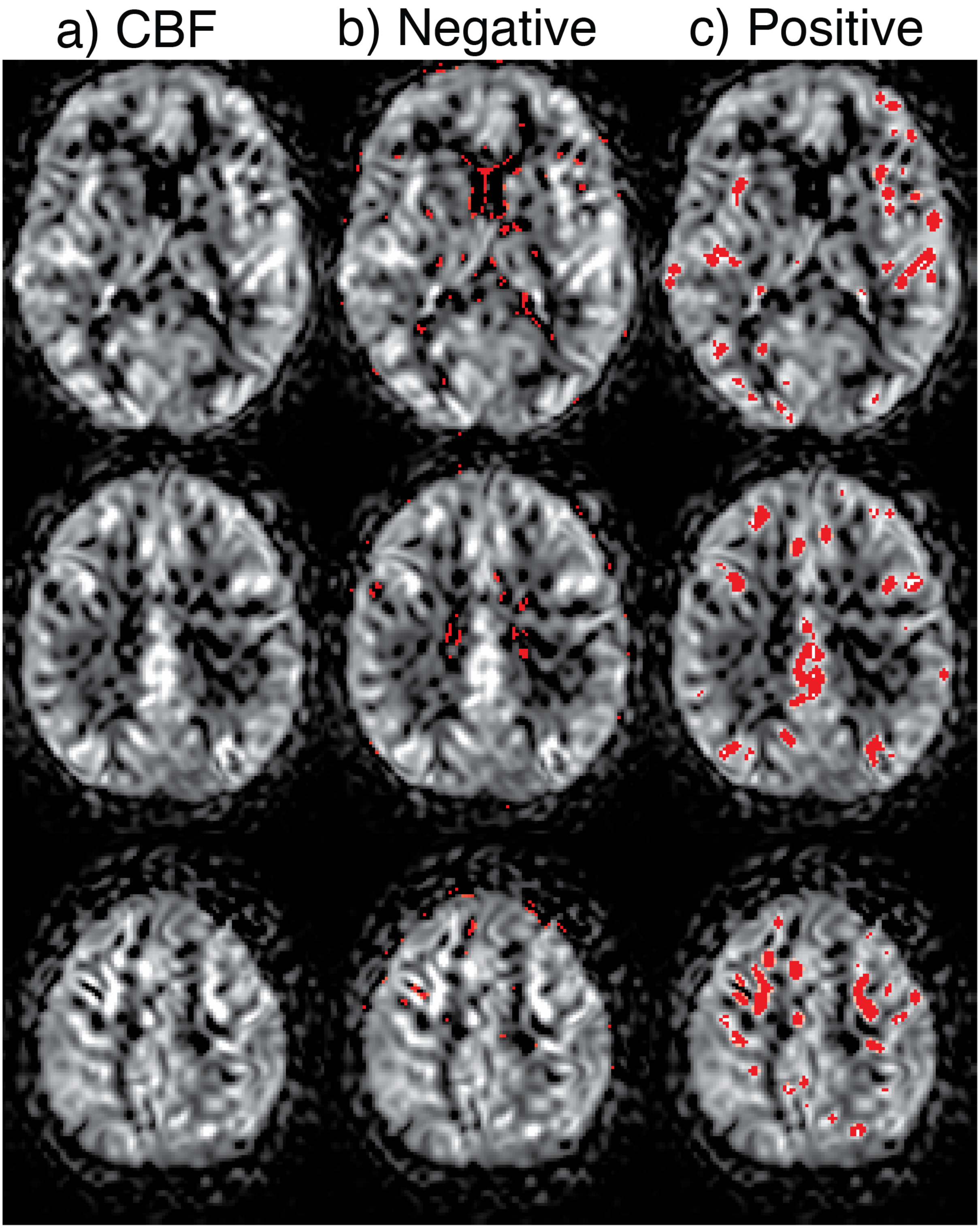


Supplementary Figure 9. Example of intra-vascular artifacts, shown without (a) and with segmentation – shown in red – of the negative cerebral blood flow (CBF) clusters (b) and extreme positive CBF voxels (c). First, spatially connected subzero voxels are grouped into clusters. The average CBF of each cluster is obtained, and clusters with significant negative mean CBF are isolated (median - 3 median absolute difference (MAD) within all subzero CBF voxels) (b) (Maumet et al., 2012). Likewise, voxels with extreme positive signal are detected by having intensity of more than median + 3 MAD within all positive CBF voxels (c). Data was from the same EPAD participant as Supplementary Figures 5 and 6 (Ritchie et al., 2016; Ten Kate et al., 2018).

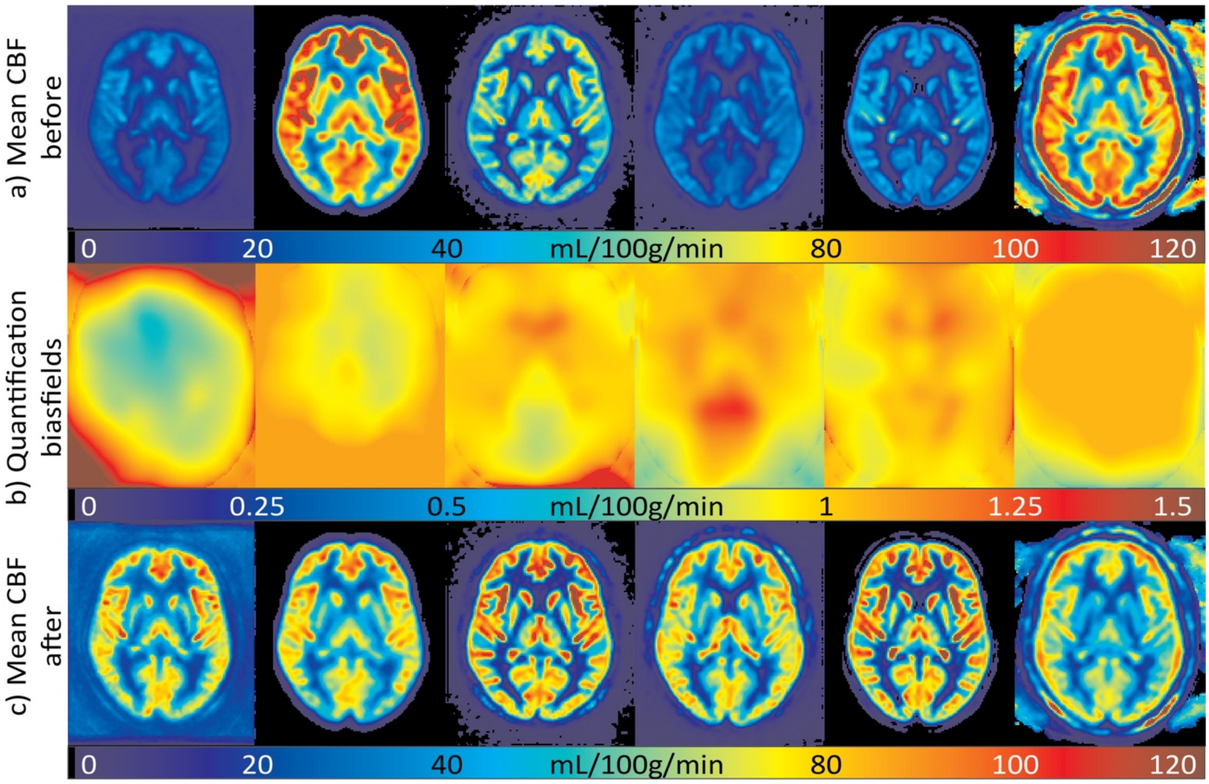


Supplementary Figure 10. An example of the intensity-normalization procedure for the six datasets shown in Figure 2. The mean CFB is shown before (a) and after (c) application of the intensity biasfield (b) that corrects for global and regional scaling between the sequences (Mutsaerts et al., 2019).

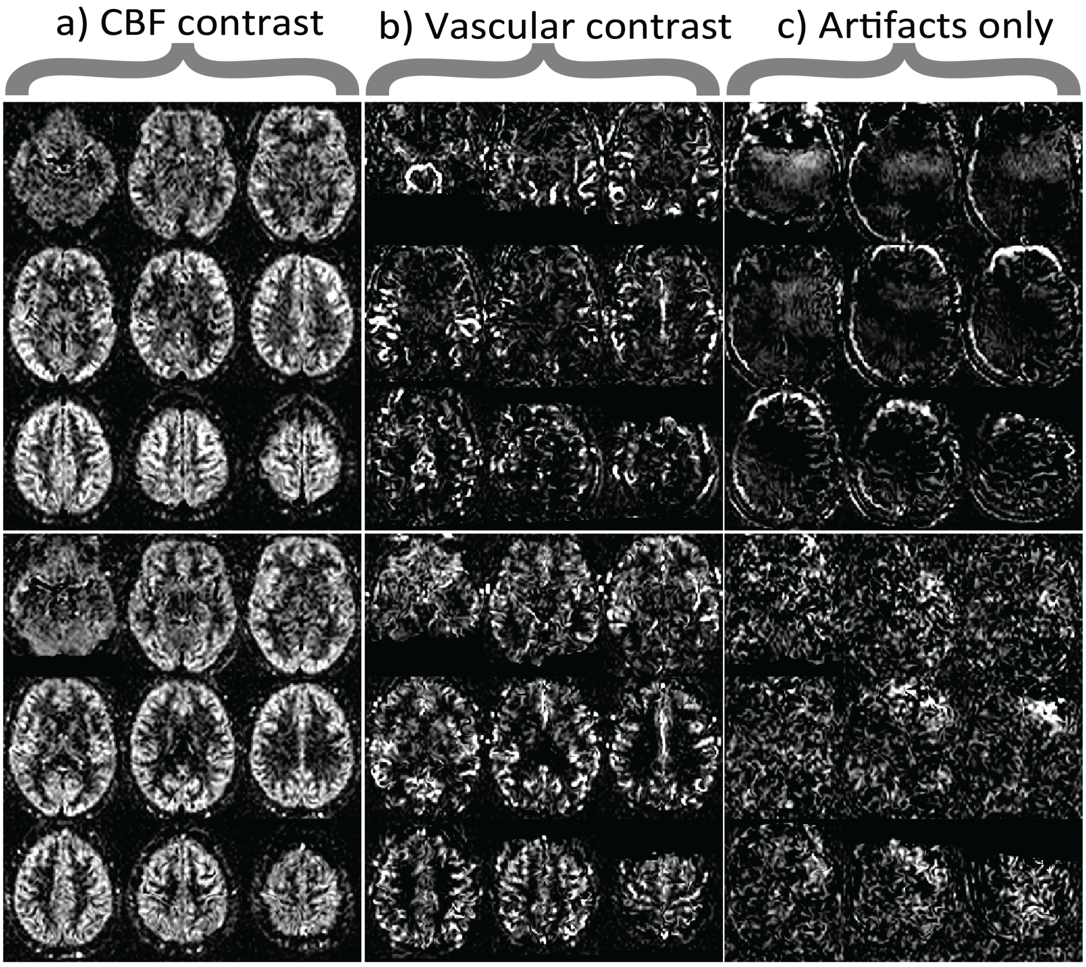


Supplementary Figure 11. Examples of CBF quality control JPG image files, allowing easy and quick browsing across the whole population for fast exploration. The CBF images can be categorized into a) CBF contrast, b) vascular contrast and c) no CBF or vascular contrast (Mutsaerts et al., 2019). CBF = cerebral blood flow, SNR = signal-to-noise ratio. Data from the Singapore Memory Clinic Study (Ferro et al., 2019).

### References

Alsop, D.C., Detre, J.A., Golay, X., Günther, M., Hendrikse, J., Hernandez-Garcia, L., Lu, H., MacIntosh, B.J., Parkes, L.M., Smits, M., van Osch, M.J.P., Wang, D.J.J., Wong, E.C., Zaharchuk, G., 2015. Recommended implementation of arterial spin-labeled perfusion MRI for clinical applications: A consensus of the ISMRM perfusion study group and the European consortium for ASL in dementia. Magn. Reson. Med. 73, 102–116.

Atkinson, D., Hill, D.L., Stoyle, P.N., Summers, P.E., Keevil, S.F., 1997. Automatic correction of motion artifacts in magnetic resonance images using an entropy focus criterion. IEEE Trans. Med. Imaging 16, 903–910.

Baas, K.P.A., Mutsaerts, H., Petr, J., Kuijer, J.P.A., Van de Ven, K., 2018. Comparing pCASL CBF measurements between 3D-GraSE and 2D-EPI on 1.5T and 3T systems, in: Proc. Intl. Soc. Mag. Reson. Med. 26 (2018). Presented at the ISMRM, p. 2158.

Bjørnebekk, A., Walhovd, K.B., Jørstad, M.L., Due-Tønnessen, P., Hullstein, I.R., Fjell, A.M., 2017. Structural Brain Imaging of Long-Term Anabolic-Androgenic Steroid Users and Nonusing Weightlifters. Biol. Psychiatry 82, 294–302.

Blokhuis, C., Mutsaerts, H.J.M.M., Cohen, S., Scherpbier, H.J., Caan, M.W.A., Majoie, C.B.L.M., Kuijpers, T.W., Reiss, P., Wit, F.W.N.M., Pajkrt, D., 2017. Higher subcortical and white matter cerebral blood flow in perinatally HIV-infected children. Medicine 96, e5891.

Clement P, Vanlessen N, Mutsaerts HJMM, Bogaert S, Pourtois G, Achten E, n.d. Mood-related changes of cerebral perfusion in hu mans using arterial spin labeling, in: ESMRMB 2016. Presented at the European Society of Magnetic Resonance in Medicine and Biology.

Dahnke, R., Ziegler, G., Großkreutz, J., Gaser, C., 2015. Quality Assurance in Structural MRI, in: HBM. Honolulu.

Elvsåshagen, T., Mutsaerts, H.J., Zak, N., Norbom, L.B., Quraishi, S.H., Pedersen, P.Ø., Malt, U.F., Westlye, L.T., van Someren, E.J., Bjørnerud, A., Groote, I.R., 2018. Cerebral blood flow changes after a day of wake, sleep, and sleep deprivation. Neuroimage. https://doi.org/10.1016/j.neuroimage.2018.11.032

Ferro, D.A., Mutsaerts, H.J., Hilal, S., Kuijf, H.J., Petersen, E.T., Petr, J., van Veluw, S.J., Venketasubramanian, N., Boon Yeow, T., Jan Biessels, G., Chen, C., 2019. Cortical microinfarcts in memory clinic patients are associated with reduced cerebral perfusion. J. Cereb. Blood Flow Metab. 271678X19877403.

Gaser, C., 2009. Partial Volume Segmentation with Adaptive Maximum A Posteriori (MAP) Approach. Neuroimage 47, S121.

Kurth, F., Gaser, C., Luders, E., 2015. A 12-step user guide for analyzing voxel-wise gray matter asymmetries in statistical parametric mapping (SPM). Nat. Protoc. 10, 293–304.

Liu, T.T., 2017. Reprint of “Noise contributions to the fMRI signal: An Overview.” Neuroimage 154, 4–14.

Magnotta, V.A., Friedman, L., FIRST BIRN, 2006. Measurement of Signal-to-Noise and Contrast-to-Noise in the fBIRN Multicenter Imaging Study. J. Digit. Imaging 19, 140–147.

Maumet, C., Maurel, P., Ferré, J.-C., Bannier, E., Barillot, C., 2012. Using negative signal in mono-TI pulsed arterial spin labeling to outline pathological increases in arterial transit

times. ISMRM Scientific Workshop. Perfusion MRI: Standardization, Beyond CBF & Everyday Clinical Applications 40, 42.
Mutsaerts, H.J.M.M., Mirza, S.S., Petr, J., Thomas, D.L., Cash, D.M., Bocchetta, M., de Vita, E., Metcalfe, A.W.S., Shirzadi, Z., Robertson, A.D., Tartaglia, M.C., Mitchell, S.B., Black, S.E., Freedman, M., Tang-Wai, D., Keren, R., Rogaeva, E., van Swieten, J., Laforce, R., Tagliavini, F., Borroni, B., Galimberti, D., Rowe, J.B., Graff, C., Frisoni, G.B., Finger, E., Sorbi, S., de Mendonça, A., Rohrer, J.D., MacIntosh, B.J., Masellis, M., GENetic Frontotemporal dementia Initiative (GENFI), 2019. Cerebral perfusion changes in presymptomatic genetic frontotemporal dementia: a GENFI study. Brain 142, 1108–1120.
Mutsaerts, H.J.M.M., Richard, E., Heijtel, D.F.R., van Osch, M.J.P., Majoie, C.B.L.M., Nederveen, A.J., 2014a. Gray matter contamination in arterial spin labeling white matter perfusion measurements in patients with dementia. NeuroImage: Clinical 4, 139–144.
Mutsaerts, H.J.M.M., Steketee, R.M.E., Heijtel, D.F.R., Kuijer, J.P.A., Van Osch, M.J.P., Majoie, C.B.L.M., Smits, M., Nederveen, A.J., 2014b. Inter-vendor reproducibility of pseudo-continuous arterial spin labeling at 3 Tesla. PLoS One 9, e104108.
Mutsaerts, H., Petr, J., Thomas, D.L., De Vita, E., Cash, D.M., Van Osch, M.J.P., Golay, X., Groot, P.F.C.C., Ourselin, S., Van Swieten, J., Laforce, R., Jr, Tagliavini, F., Borroni, B., Galimberti, D., Rowe, J.B., Graff, C., Pizzini, F.B., Finger, E., Sorbi, S., Castelo Branco, M., Rohrer, J.D., Masellis, M., Macintosh, B.J., 2018. Comparison of arterial spin labeling registration strategies in the multi-center GENetic frontotemporal dementia initiative (GENFI). J. Magn. Reson. Imaging 47, 131–140.
Paloyelis, Y., Doyle, O.M., Zelaya, F.O., Maltezos, S., Williams, S.C., Fotopoulou, A., Howard, M.A., 2016. A Spatiotemporal Profile of In Vivo Cerebral Blood Flow Changes Following Intranasal Oxytocin in Humans. Biol. Psychiatry 79, 693–705.
PCP Quality Assessment Protocol [WWW Document], n.d. URL http://preprocessed-connectomes-project.org/quality-assessment-protocol/ (accessed 9.2.19).
Pernet, C., 2019. SPM U+ [WWW Document]. Open Science Framework. URL https://osf.io/wn3h8/ (accessed 4.1.19).
Power, J.D., Barnes, K.A., Snyder, A.Z., Schlaggar, B.L., Petersen, S.E., 2012. Spurious but systematic correlations in functional connectivity MRI networks arise from subject motion. Neuroimage 59, 2142–2154.
Ritchie, C.W., Molinuevo, J.L., Truyen, L., Satlin, A., Van der Geyten, S., Lovestone, S., 2016. Development of interventions for the secondary prevention of Alzheimer’s dementia: the European Prevention of Alzheimer's Dementia (EPAD) project. The Lancet Psychiatry 3, 179–186.
Shirzadi, Z., Crane, D.E., Robertson, A.D., Maralani, P.J., Aviv, R.I., Chappell, M.A., Goldstein, B.I., Black, S.E., MacIntosh, B.J., 2015. Automated removal of spurious intermediate cerebral blood flow volumes improves image quality among older patients: A clinical arterial spin labeling investigation. J. Magn. Reson. Imaging 42, 1377–1385.
Ten Kate, M., Ingala, S., Schwarz, A.J., Fox, N.C., Chételat, G., van Berckel, B.N.M., Ewers, M., Foley, C., Gispert, J.D., Hill, D., Irizarry, M.C., Lammertsma, A.A., Molinuevo, J.L., Ritchie, C., Scheltens, P., Schmidt, M.E., Visser, P.J., Waldman, A., Wardlaw, J., Haller, S., Barkhof, F., 2018. Secondary prevention of Alzheimer’s dementia: neuroimaging contributions. Alzheimers. Res. Ther. 10, 112.

Visser, M., Müller, D.M.J., van Duijn, R.J.M., Smits, M., Verburg, N., Hendriks, E.J., Nabuurs, R.J.A., Bot, J.C.J., Eijgelaar, R.S., Witte, M., van Herk, M.B., Barkhof, F., de Witt Hamer, P.C., de Munck, J.C., 2019. Inter-rater agreement in glioma segmentations on longitudinal MRI. Neuroimage Clin 22, 101727.

Wald, L.L., Polimeni, J.R., 2017. Impacting the effect of fMRI noise through hardware and acquisition choices - Implications for controlling false positive rates. Neuroimage 154, 15–22.

Wang, Z., Aguirre, G.K., Rao, H., Wang, J., Fernández-Seara, M.A., Childress, A.R., Detre, J.A., 2008. Empirical optimization of ASL data analysis using an ASL data processing toolbox: ASLtbx. Magn Reson.Imaging 26, 261–269.

Wang, Z., Bovik, A.C., Sheikh, H.R., Simoncelli, E.P., 2004. Image quality assessment: from error visibility to structural similarity. IEEE Trans. Image Process. 13, 600–612.
